# Continuous evolution of user-defined genes at 1-million-times the genomic mutation rate

**DOI:** 10.1101/2023.11.13.566922

**Authors:** Gordon Rix, Rory L. Williams, Hansen Spinner, Vincent J. Hu, Debora S. Marks, Chang C. Liu

## Abstract

When nature maintains or evolves a gene’s function over millions of years at scale, it produces a diversity of homologous sequences whose patterns of conservation and change contain rich structural, functional, and historical information about the gene. However, natural gene diversity likely excludes vast regions of functional sequence space and includes phylogenetic and evolutionary eccentricities, limiting what information we can extract. We introduce an accessible experimental approach for compressing long-term gene evolution to laboratory timescales, allowing for the direct observation of extensive adaptation and divergence followed by inference of structural, functional, and environmental constraints for any selectable gene. To enable this approach, we developed a new orthogonal DNA replication (OrthoRep) system that durably hypermutates chosen genes at a rate of >10^−4^ substitutions per base *in vivo*. When OrthoRep was used to evolve a conditionally essential maladapted enzyme, we obtained thousands of unique multi-mutation sequences with many pairs >60 amino acids apart (>15% divergence), revealing known and new factors influencing enzyme adaptation. The fitness of evolved sequences was not predictable by advanced machine learning models trained on natural variation. We suggest that OrthoRep supports the prospective and systematic discovery of constraints shaping gene evolution, uncovering of new regions in fitness landscapes, and general applications in biomolecular engineering.

## Introduction

Over the history of life, evolution has carried out a large-scale experiment exploring how gene sequences change under the constraints of prevailing or shifting structural and functional demands. The results of this natural experiment, embedded within the patterns of diversity across extant gene sequences, are of fundamental value to almost all areas of life sciences. For example, sequence conservation within a gene family is used to identify functionally critical residues^1– 4^, covariation among positions in an RNA or protein is used to deduce structural contacts and sectors of connectivity^5–12^, differences in amino acid composition reveal environmental preferences (*e*.*g*., temperature^13–15^ or subcellular localization^16^), and differences in the conserved physicochemical properties across regions of a protein reflect driving forces behind folding^17,18^. Natural diversity across homologs also serves as a shared biomolecular engineering resource that can be mined for desired activities or recombined to access new functions^19–25^. Additionally, machine learning (ML) models have proven incredibly effective at extracting meaningful representations of biomolecular structure and function from the extensive diversity within and across gene families, as exemplified by their ability to predict functional effects of mutations^26–28^, design functional sequences^29–31^, and predict protein structures^32–34^. However, the natural evolution of highly diverse gene sequences under the constraints of selective forces — or conversely, the imprinting of selective forces and design principles into the statistics of sequence diversity — takes a long time at the slow rates of mutation in cellular and multicellular organisms. For example, reaching the 11% median divergence separating essential mouse and human genes^35^ took ∼96 million years^36^. Moreover, generating extensive collections of diverged sequences required complex histories of geographical isolation and speciation that allowed many populations to evolve separately to maintain variation in the face of within-population selective sweeps or genetic drift. Is it possible to compress long and vast gene evolution processes into laboratory experiments? Doing so would allow us to systematically detect novel structural and functional constraints governing biology, engineer custom biomolecules, create rich new sources of genetic variation for neofunctionalization or ML, and prospectively study the mechanisms and principles by which histories of selective forces become embedded into the patterns of sequence diversity.

We and others have endeavored towards this goal^37,38^ through the development of scalable accelerated continuous evolution systems^39,40^, such as our orthogonal DNA replication (OrthoRep) system in *Saccharomyces cerevisiae*^41,42^. OrthoRep cells have an additional DNA replication system comprising an orthogonal DNA polymerase (DNAP)/plasmid pair wherein the orthogonal DNAP (TP-DNAP1) durably replicates the orthogonal plasmid (p1) but does not replicate the host genome; likewise, host DNAPs replicate the host genome but not p1 (Fig. 1A). Through this architecture, OrthoRep supports the sustainable coexistence of two independent mutation rates in the same cell: a low mutation rate of 10^−10^ substitutions per base (s.p.b.) for the large host genome, and a high mutation rate of 10^−5^ s.p.b. exclusively acting on p1 encoding only user-defined genes. OrthoRep’s 10^−5^ s.p.b. mutation rate exceeds the error thresholds of the host genome^42^, allowing us to drive the rapid, continuous evolution of chosen genes as cells autonomously propagate. However, 10^−5^ s.p.b. is not high enough to observe extensive evolution on laboratory timescales in the general case where evolution occurs both with and without positive selection. In the specific case of evolution under strong positive selection, sufficiently large population sizes can directly compensate for moderate mutation rates by increasing the beneficial mutation supply on which selection “pulls”; in this case, the OrthoRep system has successfully evolved enzymes^38,43,44^, biosynthetic pathways^45^, biosensors^46^, drug targets^42^, and antibodies^47,48^ through long adaptive mutational pathways. Yet in the general case that includes when purifying selection is dominant or when selection is absent, both highly relevant in the generation of natural diversity and the ability to escape local optima, mutation becomes the main force pushing sequence change. Without the pull of positive selection, our previous OrthoRep systems would take 100 generations (8-12 days for the yeast host of OrthoRep) just to sample an average of 1 new mutation in a typical 1 kb gene.

**Figure 1.**
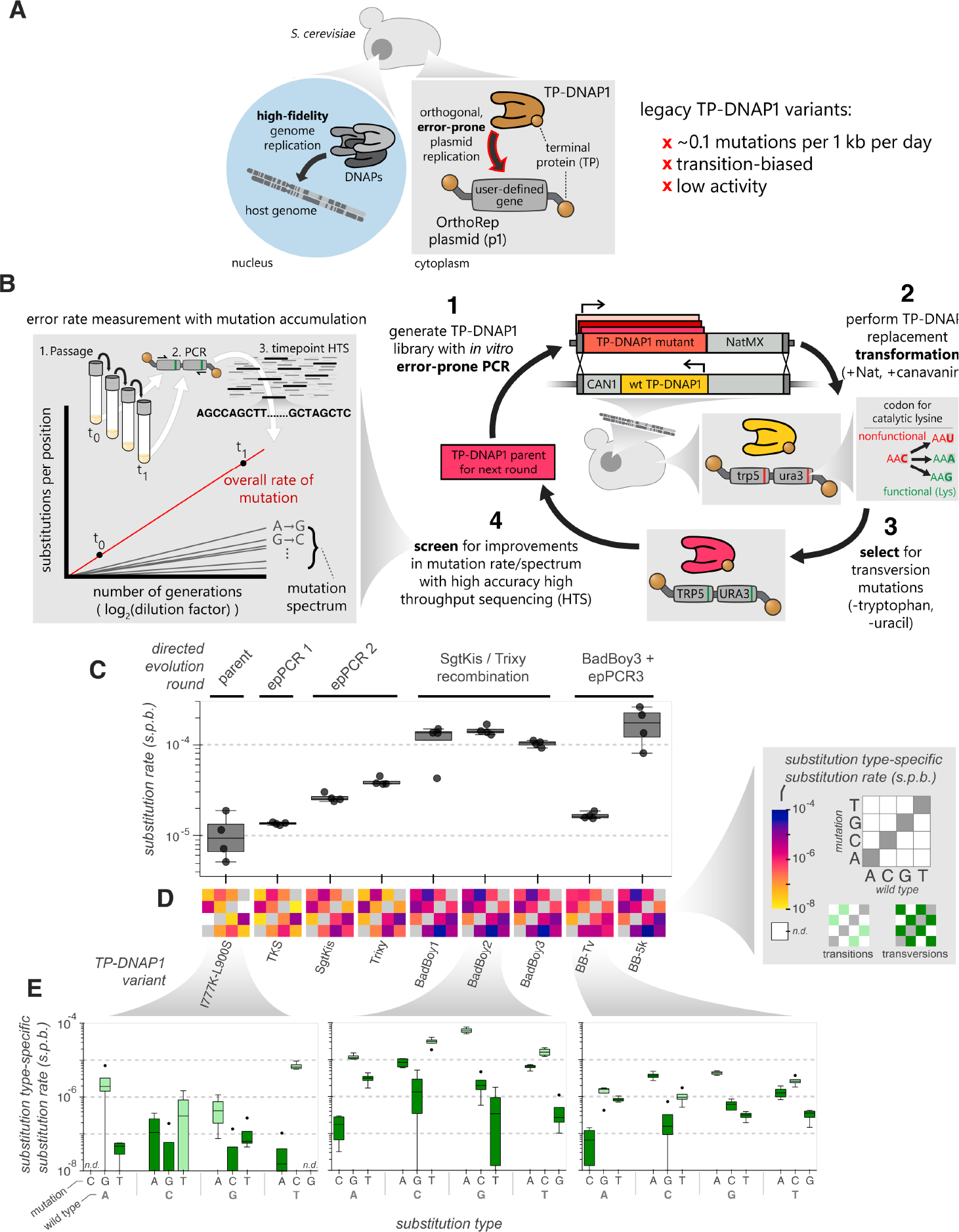
Engineering orthogonal DNA polymerases for increased mutation rates. (**A**) Architecture of the OrthoRep system. A DNA polymerase (TP-DNAP1) that exclusively replicates a specific cytoplasmically localized plasmid via protein primed replication at a high error rate enables *in vivo* targeted mutagenesis without mutagenizing genomic DNA. (**B**) Schematic for a directed evolution approach to improving TP-DNAP1’s mutation rates and mutation spectrum incorporating both a direct selection for rare transversion mutations as well as high accuracy mutation rate measurement using a mutation accumulation and high throughput sequencing (HTS) assay. (**C**-**E**) Mutation rate measurements via mutation accumulation for a series of TP-DNAP1 directed evolution intermediates showing either overall mutation rates as boxplots (C), mean mutation rates for individual substitution types as heatmaps (D), or mutation rates for individual substitution types for three individual TP-DNAP1 variants as boxplots (E). Points are representative of individual biological replicates, each representing 3-4 timepoints with >50 sequences each. Boxplots and heatmaps are representative of n=4 biological replicates. Box plot central line, boxes, and whiskers represent the median, interquartile range, and minimum / maximum values, respectively.

Here, we present the engineering of novel OrthoRep systems that have a mutation rate up to 1.7 × 10^−4^ s.p.b., corresponding to a new mutation in a typical 1 kb gene once every <10 generations in the absence of any selection. This intensified mutational force on chosen genes, operating at >1 million times the mutation rate of the host genome, allows us to mimic extended periods of natural gene evolution on laboratory timescales in the general case. We describe the TP-DNAP1 engineering effort leading to these novel OrthoRep systems and provide characterization of their performance through detailed measurements of their elevated mutation rates, reduced mutational biases, and durability. We then show how, in <3 months of laboratory passaging (totaling <15 hours of researcher intervention) of 96 independent populations, a conditionally essential gene encoded on p1 diverges to an extent where the median distance separating pairs of evolved sequences is 35 amino acids, with thousands of unique pairs separated by ≥60 amino acids. This corresponds to an amino acid divergence of ∼9% to >15%, exceeding the median 11% distance separating orthologous genes between mouse and human^35^. By analyzing the rich collection of diverged sequences throughout their laboratory evolutionary history, we uncover hidden forces shaping and constraining sequence change, such as a preference for negative net charge, supporting a proposed mechanism by which proteins avoid large-scale indiscriminate clustering in the crowded environment inside cells^49^. We also extract examples of allosteric network remodeling through the cooccurrence of mutations across distinct clades and temperature optimization through amino acid content change. Overall, our work provides an approach to systematically reveal the evolutionary constraints and selective forces that genes experience and delivers an upgraded OrthoRep system for broad application.

## Results

### Motivation and preparation for OrthoRep engineering

The current state-of-the-art OrthoRep system uses TP-DNAP1-4-2 as the error-prone orthogonal DNAP^42^. Besides its suboptimal error rate of 10^−5^ s.p.b., TP-DNAP1-4-2 also has low replicative activity (Fig. S1) and exhibits a heavily transition-biased mutation spectrum (Fig. S2)^42^, suppressing the impact of point mutations on amino acid sequence during protein evolution (Fig. S3). To increase OrthoRep’s overall error rate, transversion rate, and activity, we carried out a directed evolution campaign on TP-DNAP1.

To prepare the directed evolution campaign, we engineered a genetic selection strain, OR-Y488, that could enrich TP-DNAP1s with increased error rates and activities from libraries of TP-DNAP1 variants (Fig. 1B). We prioritized selection for increased transversion rates under the rationale that TP-DNAP1s with high transversion rates would also have high overall error rates, because transversions require tolerance of larger structural aberrations than transitions^50^. OR-Y488 contained a p1 plasmid (p1-ura3*-trp5*) encoding two auxotrophic marker genes, *ura3* and *trp5*, each specifically disabled via an active site missense mutation whose sole option for functional reversion is a transversion (Fig. 1B and Fig. S4). TP-DNAP1s with the highest transversion rates should restore *URA3* and *TRP5* most frequently, resulting in their enrichment when OR-Y488 is grown in the absence of exogenous uracil or tryptophan. This selection was designed to occur in two sequential stages, first for *URA3* restoration and then for *TRP5* restoration, to suppress the enrichment of low error rate TP-DNAP1 variants in revertants that stochastically emerge from the long tail of the Luria-Delbrück distribution^51^. We also designed a counterselection-based high-efficiency integration strategy for transforming TP-DNAP1 variants into OR-Y488 such that only the variant TP-DNAP1 would replicate p1-ura3*-trp5* in transformed cells (see Methods and Fig. S5).

To precisely guide the TP-DNAP1 directed evolution campaign, we developed a mutation accumulation assay^52^ for p1 and coupled it with high throughput sequencing (HTS) of p1 amplicons using the Oxford Nanopore Technologies (ONT) platform^53^, allowing us to accurately determine the rate for any individual type of mutation (Fig. 1B). In this assay, a strain containing p1 replicated by any TP-DNAP1 variant is grown for a set number of generations. A region of p1 not under selection is then amplified, tagged with unique molecular identifiers (UMIs) for sequencing error correction^54,55^, and sequenced at two or more timepoints during mutation accumulation (see Methods). The rate of change in the number of mutations per position is calculated for all types of mutations to fully describe the overall mutation rate and mutation preferences of the TP-DNAP1 variant. To facilitate rapid characterization, we developed a custom analysis pipeline that could carry out most of the analysis steps autonomously (Fig. S6). This pipeline, <u>M</u>utation <u>A</u>nalysis for <u>P</u>arallel <u>L</u>aboratory <u>E</u>volution or Maple, performs consensus sequence generation, demultiplexing, mutation identification, mutation rate analysis, and many other operations to generate a collection of visualizations and data tables that accelerate analysis of mutation-rich sequencing datasets while minimizing user input.

### Directed evolution of a high error rate orthogonal DNAP

With the genetic selection, HTS-based mutation accumulation measurement pipeline, and Maple in place, we carried out our TP-DNAP1 directed evolution campaign over five rounds, including three rounds of error-prone PCR (epPCR) and selection (Table S1). In round 1, we started from an epPCR library generated from TP-DNAP1(I777K, L900S). TP-DNAP1(I777K, L900S) is a relative of TP-DNAP1-4-2 with a mutation rate near 10^−5^ s.p.b. and greater activity than TP-DNAP1-4-2^42^ (Fig. S1). We reasoned that its higher activity would confer mutational robustness, increasing the fraction of active library members when used as the parental sequence. After integration of the epPCR library into OR-Y488 and enrichment of TP-DNAP1s that could restore *URA3* via a transversion mutation (Fig. S4C), we isolated ∼90 colonies and screened for their ability to restore *TRP5* through the second selection stage, which was done in multiple replicates per clone following a fluctuation analysis format^42^ to obtain estimates on phenotypic mutation rate. We isolated several clones whose phenotypic mutation rates were up to 10-fold elevated (Figs. S7A) and evaluated their genotypic mutation rates in detail using our mutation accumulation assay and Maple. Among these clones, TP-DNAP1-TKS, with the mutation P680T, had the highest per base mutation rate (Fig. S7B-C). In round 2, selection applied to an epPCR library generated from TP-DNAP1-TKS resulted in the enrichment of several clones whose full mutation rates and spectra were then determined (Fig. S8). These several clones turned out to represent two unique TP-DNAP1 variants. The two variants satisfied the requirements of selection via distinct strategies. One variant, TP-DNAP1-SgtKis, contained 5 non-synonymous mutations in addition to those in the parent TP-DNAP1-TKS and demonstrated an altered mutation spectrum favoring transversions but only a minimal apparent increase in overall mutation rate. Another variant, TP-DNAP1-Trixy contained three non-synonymous mutations and had the highest overall mutation rate measured, but only marginal changes to the mutation spectrum.

Since TP-DNAP1-SgtKis had an increased transversion rate and TP-DNAP1-Trixy had an increased overall substitution rate, we reasoned that their combination could yield orthogonal DNAPs with both high overall and transversion rates. We cloned seven new TP-DNAP1 variants where a subset of mutations from TP-DNAP1-SgtKis were added to TP-DNAP1-Trixy and obtained their fully described mutation rates (SgtKis / Trixy recombination round, Fig. S9). Remarkably, all TP-DNAP1 variants that included the mutations L474S and E488G from TP-DNAP1-SgtKis exhibited a dramatic elevation in their overall mutation rate, in each case bringing the per base rate to ∼10^−4^ s.p.b. (Fig. 1C, Fig. S9C), 1-million-fold higher than the yeast genomic mutation rate^42,56^. Furthermore, the broad mutation spectrum of TP-DNAP1-SgtKis was preserved in these variants (Fig. 1D-E, Fig. S9D), which we named BadBoy1, BadBoy2, and BadBoy3 (Table S1) to recognize their poor fidelity.

Our directed evolution campaign yielded two additional notable TP-DNAP1 variants resulting from combining mutations enriched from an epPCR library derived from TP-DNAP1-Trixy (epPCR 3 round, Fig. S10) with those in BadBoy3 (BadBoy3 + epPCR 3 round, Table S1). One of the resulting TP-DNAP1s, named BB-5k, exhibited a further increase in mutation rate, to 1.7 × 10^−4^ s.p.b. (Fig. 1C), representing ∼1 mutation every time a 5 kb recombinant p1 plasmid is replicated.

However, BB-5k did not durably maintain p1-ura3*-trp5* in two of four biological replicates over the ∼120 generations of mutation accumulation tested, possibly because it exerts an excessive mutational load on the *LEU2* marker used to maintain p1-ura3*-trp5*. The other DNAP of potential value, BB-Tv, has a relatively low mutation rate (1.6 × 10^−5^ s.p.b.), like that of our previous OrthoRep systems. Yet unlike previous systems, BB-Tv demonstrated a near-ideal mutation spectrum, with transversions accounting for of all mutations, up from only 2.5% for TP-DNAP1(I777K, L9000S) (Table S1, Fig. 1D). BB-Tv should therefore be useful in continuous evolution experiments involving larger targets that have lower error thresholds.

Detailed characterization of mutation accumulation across our several TP-DNAP1 variants revealed interesting trends. For example, when we compared mutation rates across different regions of p1, we found that mutation frequencies were consistently high in regions of p1 not under selection but were lower in regions encoding genes under selection (Fig. S11A-B), an effect that was most pronounced for the highest mutation rate TP-DNAP1 variants (R^2^ ≈ 0.5, Fig. S11C). This implies that the 10^−4^ s.p.b. mutation rates of our TP-DNAP1s have entered a regime in which, without purifying selection, the function of a gene is quickly degraded, suggestive of nearing gene error thresholds. We also found that the mutation rate of TP-DNAP1-Trixy was correlated with p1 length while TP-DNAP1-SgtKis did not exhibit this trend, suggesting an interplay between mutation rate and the number of bases replicated that is dependent on mutation mechanism (Fig. S12). For BadBoy1, BadBoy2, and BadBoy3, mutation rates were largely independent of p1 length (Fig. S12). Finally, examination of substitution-type-specific mutation rates among our TP-DNAP1 variants showed that mutation of position 777 to either Ser or Thr yields a large drop (∼10-fold) in the A:T→G:C transition mutation rate while having little effect on the reverse G:C→A:T rate (Fig. S13), demonstrating that even similar mutation types can be generated by independent mechanisms. These subtle observations that come through our precise and rigorous mutation rate measurement pipeline should aid the future engineering of OrthoRep and other continuous evolution systems^57^.

Overall, our orthogonal TP-DNAP1 directed evolution campaign, in conjunction with past efforts, complete a set of OrthoRep systems evenly spanning a range of ∼5 orders-of-magnitude, from ∼10^−9^ s.p.b., similar to the mutation rate of modern cellular genomes, up to ∼10^−4^ s.p.b., far beyond the error thresholds of modern cellular genomes^42^ and likely approaching the error threshold of individual genes where maximal adaptation rates can be reached^58,59^.

### Extensive divergence of a conditionally essential gene on laboratory timescales

Our new OrthoRep systems should be capable of driving rapid evolution of chosen genes regardless of the type of selection imposed. We encoded the β-subunit of *Thermotoga maritima’s* tryptophan synthase (TrpB) onto p1 in a tryptophan auxotroph where p1 is exclusively replicated by BadBoy2 at a mutation rate of 1.4 × 10^−4^ s.p.b. (Fig. 2A). TrpB condenses indole with serine to yield tryptophan (Trp), but *T. maritima* TrpB is maladapted for this standalone reaction, since it normally functions in complex with TrpA^60,61^. Therefore, cells grown in the absence of Trp need to evolve improved TrpB activity to propagate, allowing TrpB to serve as the subject of an extended evolution experiment that included no selection, positive selection, and purifying selection phases (Fig. S14).

**Figure 2.**
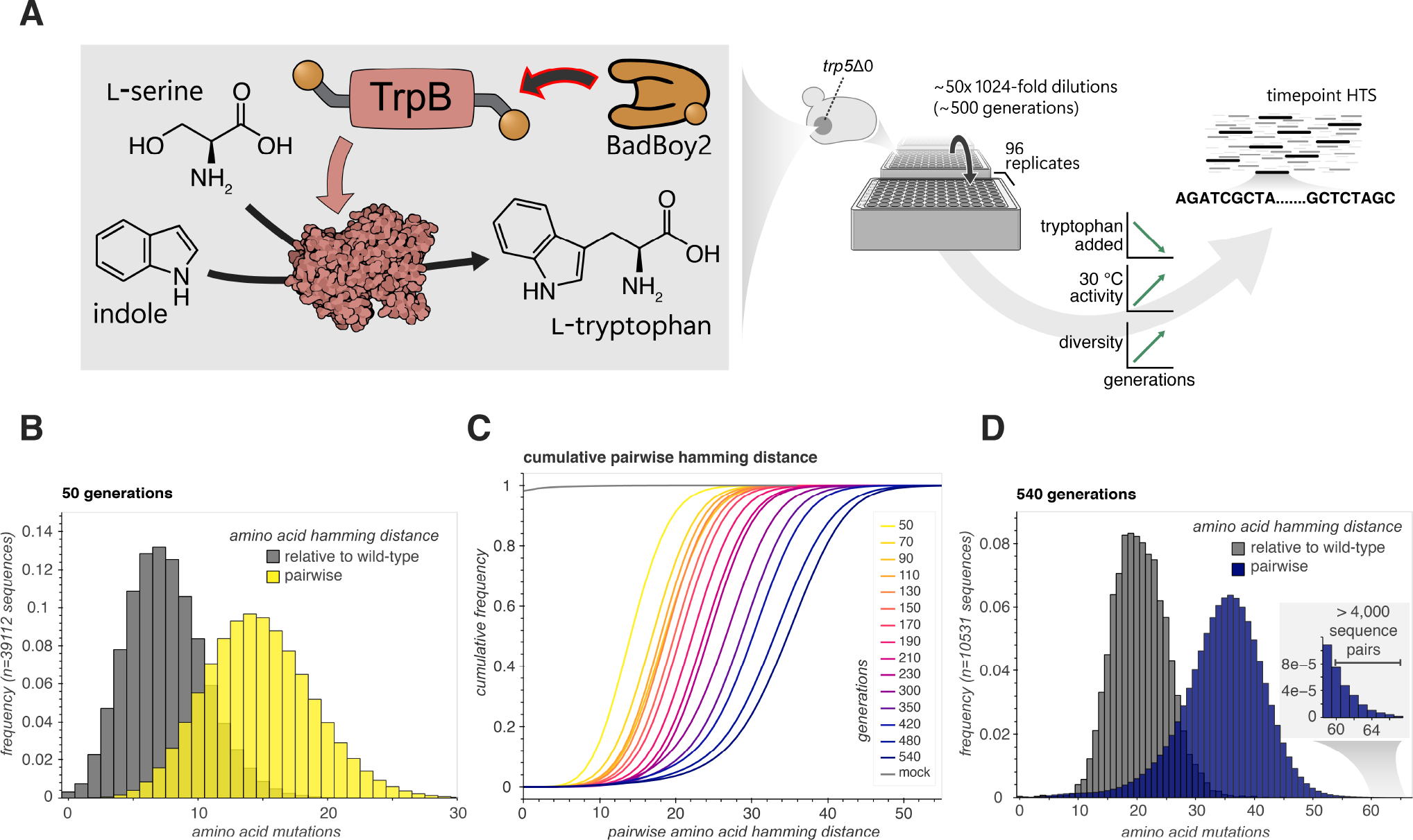
Massively parallel diversification and evolution of TrpB. (**A**) Schematic for OrthoRep evolution of the tryptophan synthase β-subunit from *Thermotoga maritima* (TrpB) for standalone function in yeast. TrpB was integrated onto the p1 plasmid in a yeast strain lacking the native yeast tryptophan synthase gene (*TRP5*). 96 independent cultures of the resulting strain were passaged mostly under selective pressure for Trp production using exogenously supplied indole over ∼500 generations. DNA from fifteen timepoints throughout the evolution campaign was harvested and sequenced using HTS. (**B**-**D**) Pairwise amino acid hamming distances as distributions for both pairwise and relative to the wild-type sequence at the first and last timepoint (B and D respectively), and as pairwise cumulative distributions for all timepoints (C).

We designed our evolution experiment to prioritize sequence divergence and diversity in order to maximize the amount of evolutionary information that could later be extracted. The evolution experiment was therefore run for ∼540 generations (<3 months) at the scale of 96 independent replicate 500 μL cultures. 1:1024 (10 generation) transfers into fresh growth medium were made every one or two days, depending on cell density, following a passaging schedule that included all types of selection pressures: ‘no selection’ phases, ‘positive selection’ phases, and ‘purifying selection’ phases, as summarized in Fig. S14. We collected DNA from cells at 15 timepoints throughout the ∼540-generation evolution experiment and used a high-yield rolling circle amplification-based sequencing strategy in conjunction with HTS and Maple to analyze the TrpB sequences sampled from these timepoints (See Methods and Fig. S15).

Overall, we observed a wide diversity of evolutionary outcomes (Fig. S16) and a monotonic increase in both the average number of mutations and diversity (as measured by pairwise hamming distances) in TrpB throughout all phases of evolution (Fig. 2B-D). The rate at which mutations accumulated in the population was highest in the no selection phase (∼0.15 amino acid changes per generation) followed by the positive selection and purifying selection phases, which exhibited similar rates (∼0.024 and ∼0.018 amino acid changes per generation, respectively) (Table S2). Notably, the positive selection phases did not have the highest rate of mutation accumulation even though there was substantial adaptation in producing TrpBs that supported cell growth in the absence of Trp and presence of moderate concentrations of indole. This suggests that BadBoy2’s error rate was high enough to consistently “saturate” positive selection with an overabundance of beneficial mutations in TrpB, predicting the broader power of upgraded OrthoRep mutation rates in biomolecular evolution applications. Notably, mutation accumulation in the purifying selection phases was appreciable yet substantially slower than in the no selection and positive selection phases. From this, we conclude that BadBoy2’s error rate was high enough to constantly test the constraints of structure and function, demonstrating the general utility of OrthoRep in uncovering biological forces governing how genes and biomolecules operate. At the end of the evolution experiment, each TrpB sequence had an average of 20.6 amino acid and 44.5 nucleotide mutations (Table S2).

In the last two timepoints, over 1800 sequences (∼5%) had accumulated more than 30 amino acid changes from the ancestral 398 amino acid wt TrpB. The distribution of pairwise hamming distances showed that sequences had substantially diverged from each other (Fig. 2C-D) such that in the final timepoint, 24% of sequences were separated from each other by 40 amino acids or more, including more than 4000 sequence pairs differing by a pairwise amino acid hamming distance of at least 60 (Fig. 2D, inset). This level of sequence divergence (>15%) approximates that between human and mouse essential gene orthologs^35^. Indeed, our experiment demonstrates that we can compress extensive and complex gene evolution processes into laboratory timescales, resulting in the generation of highly diverse sequence families shaped by varied selection conditions over long mutational pathways. What does the evolutionary information embedded into this diversity reveal?

### General structural and functional constraints

∼500,000 sequences of TrpB with an average of 13.1 amino acid replacements each were captured over the evolution experiment, and over 90% of those sequences were unique (Table S2). With such a diverse evolutionary dataset, patterns of conservation should contain structural and functional constraints defining TrpB. To test this notion, we used an AlphaFold structure^33^ and knowledge from previous studies on TrpB^62,63^ to first categorize each residue in TrpB according to its general structural or functional role, as outlined in Fig. 3A. We then asked whether different categories showed different levels of conservation. We immediately noticed a congruence between relative conservation and buried residues, revealing the well-known importance of a buried hydrophobic core in protein folding (Fig. 3B). We also noticed tha t residues within 5 Å of TrpB’s active site were highly conserved. Additionally, there was a relative abundance of amino acid replacements at certain positions in the COMM domain, suggesting that it was a target of adaptation.

**Figure 3.**
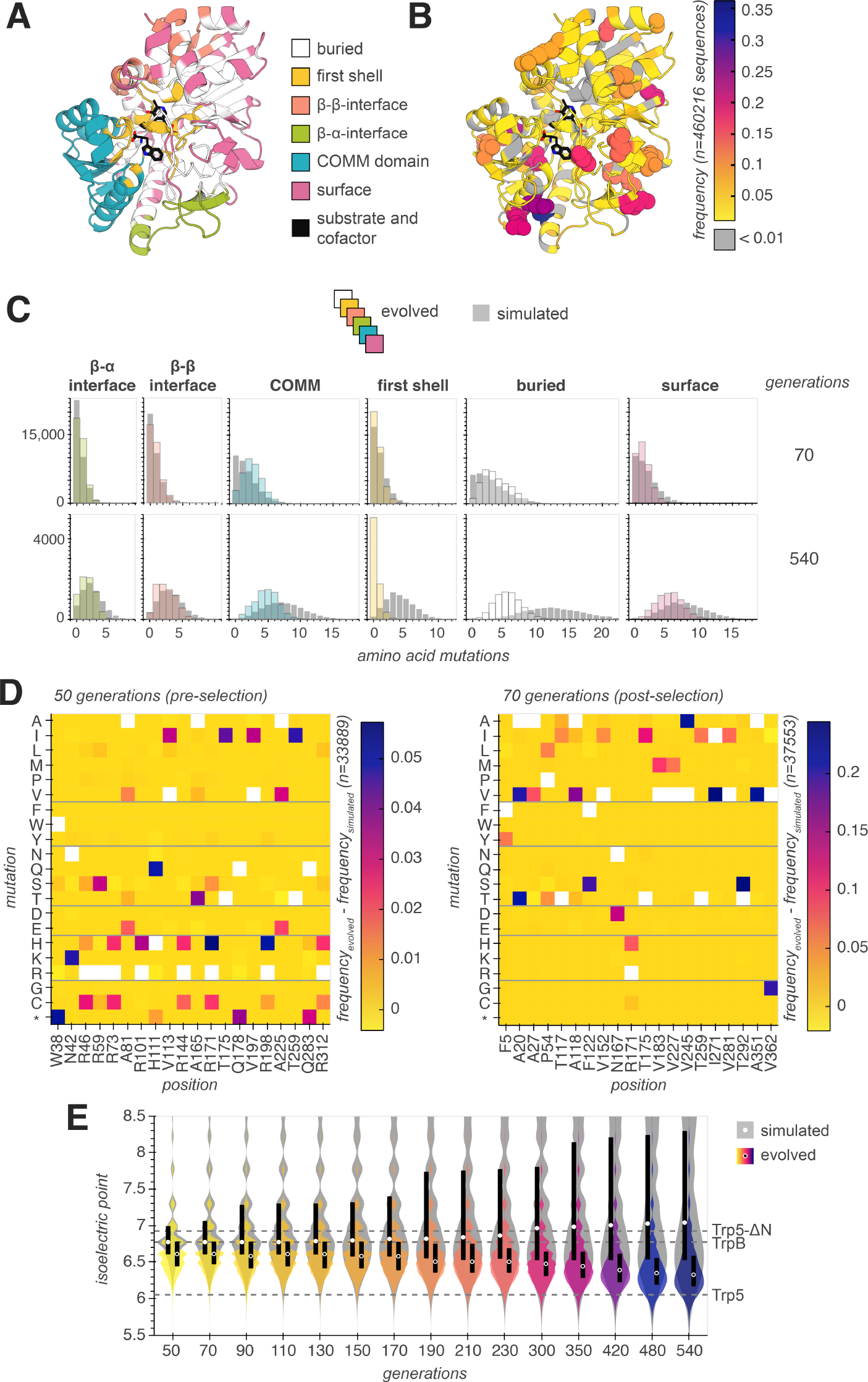
Revealed effects of selective pressures. (**A**) AlphaFold structure of *Thermotoga maritima* TrpB with different regions colored according to their structural role. First shell, β-β interface, and β-α interface residues are designated as such if they are within 5 Å of the substrate and cofactor (Trp and PLP), the other β subunit, or the other α subunit in the αββα heterotetramer holoenzyme, respectively. Alignment to *Pyrococcus furiosus* TrpB crystal structures (PDB codes 5E0K and 5DW3) were used to determine distances from substrate and cofactor, α-subunit, and β-subunit. Mean solvent accessible surface area (SASA) was used to categorize all remaining residues as either surface (SASA≥0.2) or buried (SASA<0.2). (**B**) Heatmap of mutations among OrthoRep-evolved TrpB sequences applied to the AlphaFold structure. (**C**) Distributions of mutations among OrthoRep-evolved TrpB sequences within the 6 structural regions compared to a simulated dataset of sequences with the same number of synonymous mutations. (**D**) Heatmap of mutation frequency for all mutations among the 20 most frequently mutated positions in the timepoint corresponding to either 50 or 70 generations. Frequencies for simulated sequences are subtracted to account for bias due to BadBoy2’s mutation preference and wt TrpB sequence content. Isoelectric points of the wild type TrpB, the TrpA-TrpB holoenzyme ortholog from *Saccharomyces cerevisiae* (Trp5) and an N-terminally-truncated Trp5 homologous to TrpB (Trp5-ΔN) are shown for comparison. (**E**) Violin plots of isoelectric points for all OrthoRep-evolved and simulated TrpB sequences, split by timepoint. Points and black bars denote the means and interquartile range for all sequences within each timepoint.

To improve our resolution in such observations, we generated a simulated dataset of TrpB mutants where each sequence accumulated mutations from the exact encoding of wt TrpB using BadBoy2’s fully described mutation biases. For each simulated sequence, mutation accumulation was stopped once it matched the number of synonymous mutations of a corresponding sequence from the real dataset. The simulated dataset serves as the null model where patterns in evolved TrpB sequences are simply a reflection of the mutation preferences of BadBoy2 and codon usage of wt TrpB. Under the assumption that synonymous mutations have no effect on fitness, the nonsynonymous differences between the real dataset and the simulated dataset contain the influence of selective forces. An excess of nonsynonymous changes in the real dataset compared to the simulated dataset is therefore an indication of positive selection, while the opposite signifies purifying selection. As shown in Fig. 3C, at the generation 70 timepoint, the real dataset has a paucity of nonsynonymous mutations per sequence in the active site region and buried residues and an excess of nonsynonymous mutations per sequence in the COMM domain.

Generation 70 is during the first phase of positive selection for TrpB’s operation as a standalone enzyme capable of generating tryptophan (Fig. S14), so this timepoint is most likely to reveal signatures of adaptation. (This is supported by Fig. S17, which shows the overall mutation accumulation dynamics; generation 70 is the timepoint with the greatest excess of nonsynonymous mutations relative to the simulated dataset among the timepoints for which selection for TrpB function had been applied.) At generation 70, we find that positive selection had enriched mutations to buried and COMM domain residues while purifying selection had already removed changes to the active site region. Our detection of the COMM domain as a focus of adaptation can be explained, since TrpB is a well-studied enzyme. The COMM domain mediates allosteric activation of TrpB by TrpA^60^. As our evolution experiment required TrpB to operate in a standalone manner without TrpA, the remodeling of allosteric networks through the COMM domain was an expected means to adaptation in line with previous studies of engineered TrpB standalone activity^62,63^. Had TrpB not been well-studied, the detection of this critical region for adaptation fro m the evolutionary information could have suggested such an explanation.

We also considered the final timepoint of the evolution experiment. As shown in Fig. 3C and Fig. S17, by generation 500, the influence of purifying selection had dominated, as evidenced by the paucity of mutations in the real data compared to the simulated. Although purifying selection constrained all regions of TrpB, some were clearly more constrained than others. For example, the active site region had almost no mutations (maximally 1 or 2 but mostly 0) and deviated from the simulated mutant distribution more than all other regions. Buried residues also had substantially fewer nonsynonymous mutations than the simulated dataset. In contrast, the effect of purifying selection was less pronounced on surface residues, reflecting the relative tolerance of protein surfaces to mutation. Surprisingly, this also applied to the newly solvent-exposed β-α interface region. In the absence of the α-subunit, this region should be more solvent-exposed than in TrpB’s native context. The fact that there was little noticeable difference in the effects of selection on the β-α and β-β interfaces suggests that solvent-exposure of this region had minimal impact on TrpB fitness.

### Isoelectric point evolution for intracellular compatibility

We examined the 20 residues that were mutated in greatest excess among real evolved sequences relative to simulated sequences in the first two timepoints (Fig. 3D). Despite the entire wt TrpB protein containing only 19 arginine residues in total (5%), arginine constituted 8 of these 20 most frequently mutated residues by generation 50. This came as a surprise because this enrichment occurred even before selection for TrpB function was imposed. This led us to hypothesize charge optimization as a driving selective force, because charge could influence not only TrpB function itself, but also the cellular environment within which TrpB operated. To evaluate this hypothesis, we calculated the isoelectric point (pI) of sequences throughout the evolution experiment and examined its distribution over time (Fig. 3E). Indeed, we found that the pI of sequences was significantly lower at the end of the experiment than early in the experiment (p<0.0001, Mann-Whitney U test). Comparison of pI change with simulation corroborates that this effect was driven by positive selection. A similar analysis of hydrophobicity revealed a modest decrease in hydrophobicity over time for simulated sequences that was mitigated by selection in the real data (Fig. S18). The change in hydrophobicity for the real sequences throughout the experiment was less pronounced than the change in pI, however, suggesting that charge optimization, and not polarity in general, was the dominant selective force. Notably, the majority of the shift in the pI distribution occurred in the latter half of the experiment (Fig. 3E), highlighting the importance of sustained rapid mutagenesis over long periods of evolution to embed such presumably subtle selective forces into the data.

TrpB’s pI evolved to be comfortably below the typical yeast cytosolic pH of 6.8 to 7.2 ^64^, which is consistent with the notion that intracellular proteins prefer to be negatively charged to minimize large-scale clustering with RNAs and other proteins^49,65^. One possible mechanism by which this preference could have driven the observed adaptation is through its influence on TrpB function itself, for example by increasing the diffusivity of the enzyme^66^. Another mechanism by which a preference for negative charge in TrpB could have been adaptive is by lessening its perturbation on other entities in the cell, for example by preventing spurious association or aggregation that would disturb the function of the proteome^49^. Our data does not exclude either mechanism but suggests that the latter mechanism is present. In generations 0-50 of the evolution experiment, TrpB was not under selection for function as excess Trp was supplied to the growth media. Indeed, HTS at the end of 50 generations detected the presence of many stop codons, which were mostly eliminated soon after positive selection for TrpB adaptation had been imposed (for example, HTS at the end of generation 70) (Fig. 3D). Yet at the end of 50 generations, pIs were significantly lower than that of the simulation (p<0.0001). Our observation of charge optimization even when there was no selection for the enzyme’s function demonstrates that the intracellular context can impose constraints on the physicochemical properties of proteins independent of its primary molecular function. It also highlights the value of evolving proteins *in vivo* where subtle constraints dictating intracellular compatibility can both be revealed and included in the evolutionary optimization of protein function.

### Thermoadaptation

Given that our parental *T. maritima* TrpB was from a thermophile but needed to evolve standalone activity in a mesophile, we looked for statistical evidence of thermoadaptation in our evolved sequences. Haney *et al*. studied the patterns of amino acid replacements between natural orthologous proteins in mesophilic versus thermophilic organisms and found 17 amino acid replacements that distinguished the mesophilic variants from the thermophilic variants at homologous positions in multiple sequence alignments with high confidence^13^. When we evaluated the frequency of these 17 amino acid replacements among all mutations in our evolution experiment’s outcomes, we found that replacements in the mesophilic direction were enriched (Fig. S19). As before, this illustrates the ability of extensive gene evolution to reveal selective forces through the evolutionary information embedded into the resulting diversity.

### Networks of coupled mutations

In addition to general selective forces, we investigated finer patterns in the outcomes of TrpB evolution with the expectation that these may reveal coevolving networks of amino acids responsible for adaptation. At the beginning of our evolution experiment, we had included short barcodes adjacent to the TrpB sequence integrated onto p1. This allowed us to 1) isolate the largest clades for analysis, since these should correspond to the fittest sequences, and 2) reduce the contribution of phylogeny by computationally limiting the number of sequences analyzed per clade (Fig. 4A). The latter increases the signature of mutations independently discovered across multiple clades, favoring the detection of mutations whose cooccurrence was functionally significant. Specifically, we considered clades whose barcode had at least 100 associated sequences — there were 93 such clades — and randomly downsampled to 100 sequences for clades whose members exceeded this number.

**Figure 4.**
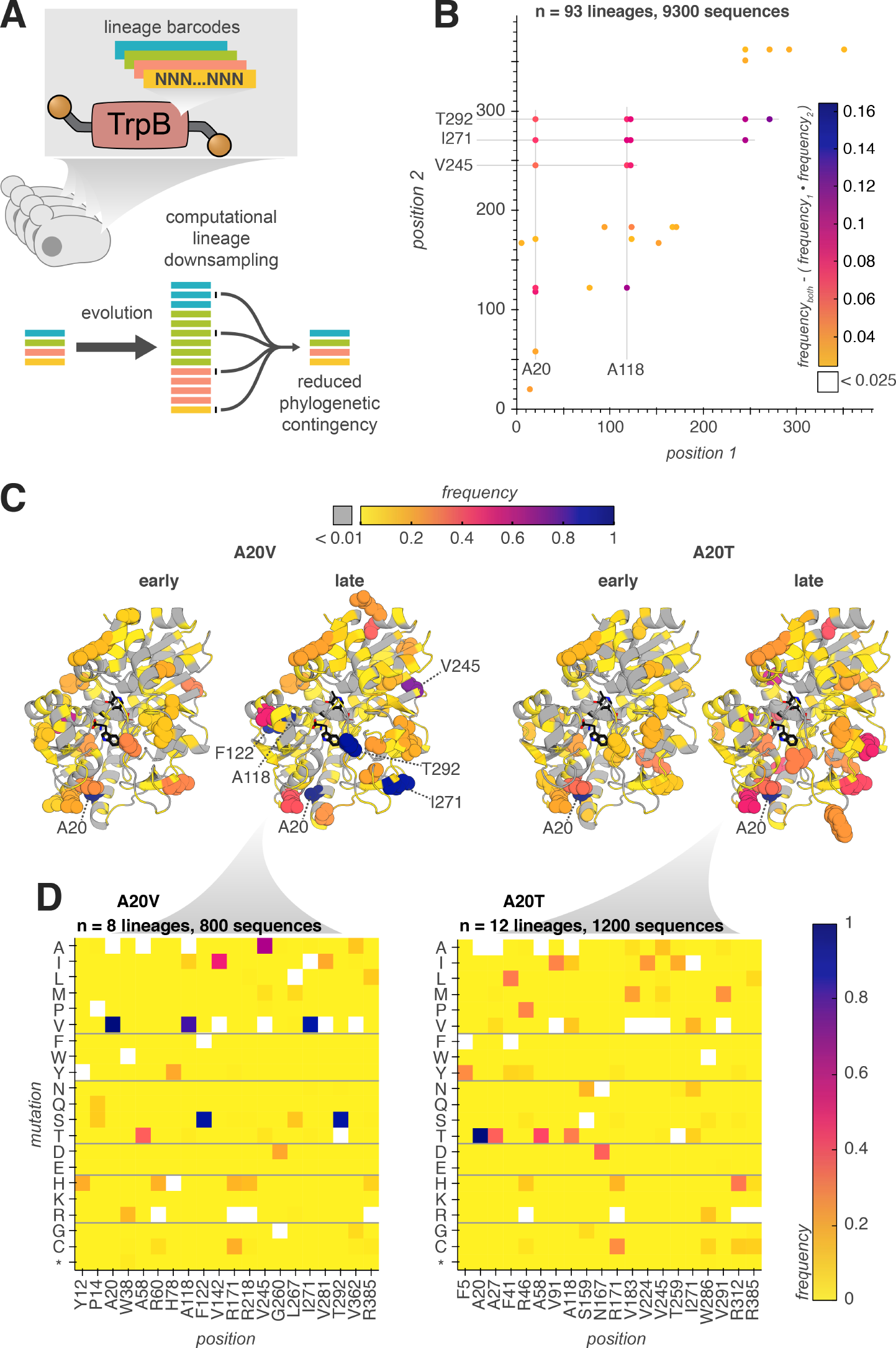
Lineage barcodes reveal covarying residues. (**A**) Schematic of computational processing used to reduce phylogenetic contingency of residue covariation. (**B**) Residue covariation plot for the most frequently covarying residues among all timepoints in the TrpB evolution dataset for 93 lineages downsampled to 100 sequences per lineage. (**C**-**D**) Heatmaps of most frequently mutated residues among sequences containing mutations A20V or A20T, downsampled to 100 sequences and chosen from specific timepoints. Heatmaps of all positions for early (generations 70 and 90) and late (generations 480 and 540) timepoints are overlaid onto a TrpB AlphaFold structure with the 20 most frequently mutated positions shown as spheres (C) or shown as mutation-specific heatmaps of the most frequently mutated 20 positions in late timepoints (D).

This analysis revealed that some sets of mutations frequently cooccur (Fig. 4B). In one particularly striking example, the A20V mutation was found to cooccur at a frequency above ∼0.7 with a set of 5 other mutations (A118V, F122S, V245A, I271V, T292S) in 8 distinct lineages in later timepoints, implying a strong relationship among these mutations (Fig. 4C-D). This set of mutations, as well as other sets identified, are spread throughout the structure of TrpB, in line with recent studies demonstrating the prevalence of structurally distributed allosterically activating mutations in TrpB and other proteins^6,62,63,67^. Among these five mutations is the T292S mutation, which was previously identified in a TrpB directed evolution campaign as highly activating, alone conferring a >5-fold increase in the standalone catalytic efficiency of TrpB^20^. Intriguingly, the association among these five mutations was sensitive to their specific identities. For example, the A20T mutation was individually present at a higher frequency than A20V among all sequences (Fig. S16D) but was not associated with any specific other mutation at a frequency above ∼0.4 (Fig. 4D). The overall picture from this analysis suggests the existence of heavily enriched individual mutations that are broadly activating across many sequence contexts and individual mutations that are only activating in combination with other mutations, implying long range epistatic interactions.

### Fitness of TrpBs and their predictability

Accurately modeling the fitness landscapes of proteins is a major goal of ML. However, it is known that ML models are biased towards favoring sequences that are more similar to the natural sequences on which they were trained^68^, and it remains unclear to what extent they can predict the function of sequences that are many mutations away from these natural sequences. It is also unclear whether ML can model how mutant sequences will perform on new functions that deviate from natural functions, in service of bioengineering goals such as enzyme and antibody engineering. To provide insight into these questions, we tested whether an advanced ML model could predict the fitness of our evolutionary outcomes.

To gain high-resolution fitness information on evolved TrpBs, we first profiled evolved variants in a high-throughput enrichment assay (Fig. 5A). We cloned a library of ∼100,000 TrpB sequences isolated from our evolution experiment into a standard yeast plasmid that would not be subject to *in vivo* mutagenesis, transformed this library into yeast, applied selection for TrpB function, and tracked the enrichment or depletion of individual variants via HTS. Included in this library were two previously engineered control TrpB variants known to be either highly functional (TrpB-003-A) or nearly nonfunctional (*Tm*Triple) in the context of yeast Trp production complementation (Fig. 5B)^38^. We evaluated Trp production by members of this library using three distinct growth conditions: Trp-supplemented media (no selection), media lacking Trp with a high concentration of indole (400 μM, weak selection), and media lacking Trp with a low concentration of indole (25 μM, strong selection). We tracked the abundance of library members in replicate yeast transformations of the same library over 4 timepoints taken at the beginning and end of 6 passages to obtain fitness scores. We found that fitness scores above a threshold (enrichment score >-5) were well correlated among replicates for both weak and strong selection conditions (R^2^=0.58 and 0.65, respectively), but not for nonselective conditions where Trp was present (Fig. 5C), confirming the reliability of the assay for highly functional TrpB variants. Thousands of multi-mutation sequences at least as functional as the previously engineered high-fitness TrpB-003-1-A, with a K_cat_/K_M_ of 1.4 × 10^5^ M^-1^s^-1^,^38^ were identified (Fig. 5D). We also observed that a large fraction of sequences had low activity (Fig. 5D), which likely owes to the multi-copy nature of p1 that creates a delay in the action of purifying selection on recently generated mutants that hitchhike with functional TrpBs in the same cell.

**Figure 5.**
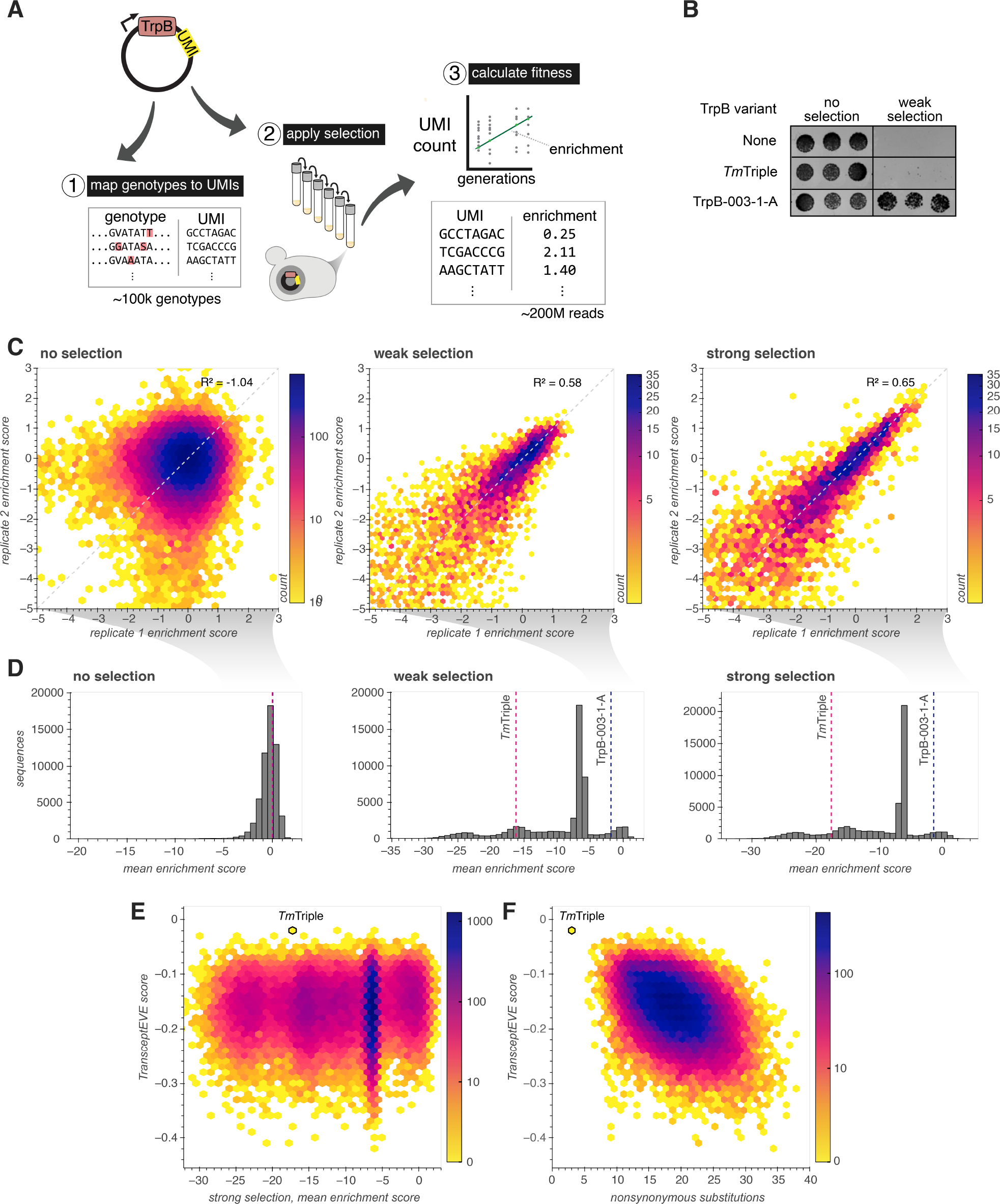
Pooled measurement and TransceptEVE prediction of TrpB variant fitness. (**A**) Schematic of pooled TrpB fitness assay using HTS. (**B**) Spot plating growth assay of control sequences included in the pooled fitness assay. (**C**) Hexbin plot of replicate concordance among pairs of replicates under growth conditions with Trp (no selection), without Trp and with 400 uM indole (weak selection) or without Trp and with 25 uM indole (strong selection) for highly functional sequences (enrichment score > -5) (**D**) Distribution of mean enrichment scores (average of n=2 biological replicates) among the three selection conditions. (**E**) Hexbin plot of TransceptEVE score vs. measured mean enrichment score with strong selection. (**F**) Hexbin plot of TransceptEVE score vs. number of nonsynonymous substitutions for all sequences with a measured strong selection mean enrichment score.

We then asked whether a state-of-the-art ML model called TranceptEVE^69^, which ensembles an autoregressive LLM (Tranception) trained across protein families with a variational autoencoder (EVE) trained on a specific family of proteins (in this case, TrpBs), could predict the measured fitness scores of our lab-evolved TrpBs. We found essentially no correlation between the relative fitness of TrpBs and their predicted fitness (Fig. 5E), even for the highly functional set of sequences (strong selection mean enrichment score >-5, Pearson correlation -0.064, p<0.0001), with the nearly nonfunctional *Tm*Triple variant being scored the highest among all sequences. In contrast and as expected, we found that scores exhibited a much stronger and negative correlation with the number of nonsynonymous amino acid mutations (Pearson correlation -0.433, p<0.0001, Fig. 5F). Our interpretation is that the evolution of TrpB for standalone function, rarely found in nature, combined with the extensiveness of sequence divergence from wt TrpB have brought our evolved sequences into regions of the fitness landscape that are out-of-distribution of natural sequences. It has been shown that large ML models can generate highly functional artificial sequences that are more dissimilar to natural sequences than our TrpBs are to wt TrpB^31^; it has also been shown that these models can nominate artificial sequences containing evolutionarily plausible mutations that improve non-natural functions^70^. Yet such ML-generated sequences are *ipso facto* within distribution of what was learned and may therefore miss important regions of underlying fitness landscapes. The ability for scalable continuous evolution to enter and explore functional regions of fitness landscapes that ML models do not, and vice-versa, highligh ts both open challenges in ML and the potential value of combining these two types of approaches going forward.

## Discussion

In this work, we have engineered OrthoRep’s mutation rate to reach >10^−4^ s.p.b. while also reducing OrthoRep’s bias against transversion mutations to maximize the exploration of sequence space. The durable action of these new mutation rates and preferences on a chosen gene, TrpB, enabled a new modality for continuous evolution where mutations quickly and durably accumulated on laboratory timescales even through periods of no selection or only purifying selection. In the absence of selection, TrpB sequences swiftly diffused into new regions of sequence space; under strong positive selection, new functional adaptations in TrpB rapidly emerged; and in the presence of purifying selection, TrpB sequences quickly sampled the space bounded by the constraints of structure and function, thereby revealing those constraints. At the end of our ∼500 generation (<3 month) evolution experiment on TrpB, we obtained thousands of unique sequences, pairs of which were separated by an average ∼35 amino acids (∼9% divergence) including many ≥60 amino acids apart (>15% divergence). The amount of evolutionary information recorded into such extensive diversity allowed us to infer both known and unknown mechanisms shaping TrpB’s evolution, including the focusing of adaptation on the COMM domain, the importance of certain allosterically linked positions on the function of TrpB, and the reduction of TrpB’s isoelectric point to yield negatively charged variants even when selection for TrpB’s enzymatic function was absent. The evolutionary information from the experiment also revealed structural and functional constraints acting on TrpB, including conservation of positions near the active site and in buried regions.

Additionally, evolved outcomes included many highly active TrpBs that exceeded the *in vivo* fitness of previously evolved and engineered TrpBs. These variants were distinct from what could be predicted by ML models trained on natural proteins, indicating the discovery of high fitness regions in the fitness landscape of TrpB that are out-of-distribution of natural variation.

While TrpB was used to demonstrate the new capabilities of OrthoRep in both the evolutionary improvement of a gene’s function and the extensive evolutionary recording of selective forces into sequence diversity, our experiments should easily extend beyond TrpB. The practicality of condensing long adaptive and neutral gene evolutionary processes into laboratory experiments should find broad applications in the evolutionary engineering of biomolecules, the mapping of sequence-function relationships, revealing novel biological constraints that shape evolution, and understanding how genes evolve — from their own points of view.

## Acknowledgements

We thank members of the Liu group for materials and thoughtful discussions. This work was funded by NIH NIGMS (R35GM136297) to CCL, NIH NCI (R01CA260415) to CCL and DSM, and a Hewitt Foundation for Medical Research Postdoctoral Fellowship to RW.

## Author contributions

Conceptualization: GR, CCL

Methodology: GR, CCL, VJH, HS

Investigation: GR, RLW, HS

Visualization: GR

Funding acquisition: CCL, DSM

Supervision: CCL, DSM

Writing – original draft: GR, CCL

Writing – review & editing: GR, CCL, RLW, HS, DSM

## Data and materials availability

All data and materials generated for this study are available upon request to the corresponding authors. The Maple sequencing analysis pipeline developed for this study is made available on Github at https://github.com/gordonrix/maple. Additional sequencing analysis scripts used for this study, experiment-specific Maple parameters, and analyzed HTS data are made available on Github at https://github.com/liusynevolab/OrthoRep_Rix_2023. Raw HTS data will be made available on NCBI’s SRA.

## Supplementary Materials

Materials and Methods

Tables S1 to S6

Figs. S1 to S19

References

## References

1. Blow, D. M., Birktoft, J. J. & Hartley, B. S. Role of a buried acid group in the mechanism of action of chymotrypsin. Nature. 221, 337–340 (1969).

2. Casari, G., Sander, C. & Valencia, A. A method to predict functional residues in proteins. Nat. Struct. Biol. 2, 171–178 (1995).

3. Capra, J. A. & Singh, M. Predicting functionally important residues from sequence conservation. Bioinformatics 23, 1875–1882 (2007).

4. Reva, B., Antipin, Y. & Sander, C. Predicting the functional impact of protein mutations: Application to cancer genomics. Nucleic Acids Res. 39, 37–43 (2011).

5. Halabi, N., Rivoire, O., Leibler, S. & Ranganathan, R. Protein Sectors: Evolutionary Units of Three-Dimensional Structure. Cell. 138, 774–786 (2009).

6. McCormick, J. W., Russo, M. A. X., Thompson, S., Blevins, A. & Reynolds, K. A. Structurally distributed surface sites tune allosteric regulation. eLife. 10, 1–38 (2021).

7. Morcos, F. et al. Direct-coupling analysis of residue coevolution captures native contacts across many protein families. Proc. Natl. Acad. Sci. U. S. A. 108, (2011).

8. Marks, D. S. et al. Protein 3D structure computed from evolutionary sequence variation. PLoS One 6, (2011).

9. Lockless, S. W. & Ranganathan, R. Evolutionarily conserved pathways of energetic connectivity in protein families. Science. 286, 295–299 (1999).

10. Rivas, E., Clements, J. & Eddy, S. R. A statistical test for conserved RNA structure shows lack of evidence for structure in lncRNAs. Nat. Methods. 14, 45–48 (2016).

11. Shindyalov, I. N., Kolchanov, N. A. & Sander, C. Can three-dimensional contacts in protein structures be predicted by analysis of correlated mutations? Protein Eng. Des. Sel. 7, 349–358 (1994).

12. Altschuh, D., Lesk, A. M., Bloomer, A. C. & Klug, A. Correlation of co-ordinated amino acid substitutions with function in viruses related to tobacco mosaic virus. J. Mol. Biol. 193, 693–707 (1987).

13. Haney, P. J. et al. Thermal adaptation analyzed by comparison of protein sequences from mesophilic and extremely thermophilic Methanococcus species. Proc. Natl. Acad. Sci. U. S. A. 96, 3578–3583 (1999).

14. Gianese, G., Argos, P. & Pascarella, S. Structural adaptation of enzymes to low temperatures. Protein Eng. 14, 141–148 (2001).

15. Berezovsky, I. N. & Shakhnovich, E. I. Physics and evolution of thermophilic adaptation. Proc. Natl. Acad. Sci. U. S. A. 102, 12742–12747 (2005).

16. Marcotte, E. M., Xenarios, I., Van Der Bliek, A. M. & Eisenberg, D. Localizing proteins in the cell from their phylogenetic profiles. Proc. Natl. Acad. Sci. U. S. A. 97, 12115–12120 (2000).

17. Shakhnovich, E., Abkevich, V. & Ptitsyn, O. Conserved residues and the mechanism of protein folding. Nature. 379, 96–98 (1996).

18. Wolfenden, R. V., Cullis, P. M. & Southgate, C. C. F. Water, Protein Folding, and the Genetic Code. Science. 206, 575–577 (1979).

19. Collias, D. & Beisel, C. L. CRISPR technologies and the search for the PAM-free nuclease. Nat. Commun. 12, 1–12 (2021).

20. Murciano-Calles, J., Romney, D. K., Brinkmann-Chen, S., Buller, A. R. & Arnold, F. H. A Panel of TrpB Biocatalysts Derived from Tryptophan Synthase through the Transfer of Mutations that Mimic Allosteric Activation. Angew. Chemie - Int. Ed. 55, 11577–11581 (2016).

21. Baier, F. et al. Cryptic genetic variation shapes the adaptive evolutionary potential of enzymes. eLife. 8, 1–20 (2019).

22. Gasiunas, G. et al. A catalogue of biochemically diverse CRISPR-Cas9 orthologs. Nat. Commun. 11, (2020).

23. Medema, M. H., de Rond, T. & Moore, B. S. Mining genomes to illuminate the specialized chemistry of life. Nat. Rev. Genet. 22, 553–571 (2021).

24. Trudeau, D. L., Smith, M. A. & Arnold, F. H. Innovation by homologous recombination. Curr. Opin. Chem. Biol. 17, 902–909 (2013).

25. Crameri, A., Raillard, S. A., Bermudez, E. & Stemmer, W. P. C. DNA shuffling of a family of genes from diverse species accelerates directed evolution. Nature. 391, 288–291 (1998).

26. Riesselman, A. J., Ingraham, J. B. & Marks, D. S. Deep generative models of genetic variation capture the effects of mutations. Nat. Methods. 15, 816–822 (2018).

27. Frazer, J. et al. Disease variant prediction with deep generative models of evolutionary data. Nature. 599, 91–95 (2021).

28. Rives, A. et al. Biological structure and function emerge from scaling unsupervised learning to 250 million protein sequences. Proc. Natl. Acad. Sci. U. S. A. 118, (2021).

29. Shin, J. E. et al. Protein design and variant prediction using autoregressive generative models. Nat. Commun. 12, 1–11 (2021).

30. Bryant, D. et al. Massively parallel deep diversification of AAV capsid proteins by machine learning. Nat. Biotechnol. 39, 691–696 (2021).

31. Madani, A. et al. Large language models generate functional protein sequences across diverse families. Nat. Biotechnol. (2023) doi:10.1038/s41587-022-01618-2.

32. Hopf, T. A. et al. Three-dimensional structures of membrane proteins from genomic sequencing. Cell. 149, 1607–1621 (2012).

33. Jumper, J. et al. Highly accurate protein structure prediction with AlphaFold. Nature. 596, 583–589 (2021).

34. Tunyasuvunakool, K. et al. Highly accurate protein structure prediction for the human proteome. Nature. 596, 590–596 (2021).

35. Makałowski, W. & Boguski, M. S. Evolutionary parameters of the transcribed mammalian genome: An analysis of 2,820 orthologous rodent and human sequences. Proc. Natl. Acad. Sci. U. S. A. 95, 9407–9412 (1998).

36. Nei, M., Xu, P. & Glazko, G. Estimation of divergence times from multiprotein sequences for a few mammalian species and several distantly related organisms. Proc. Natl. Acad. Sci. U. S. A. 98, 2497–2502 (2001).

37. Stiffler, M. A. et al. Protein Structure from Experimental Evolution. Cell Syst. 10, 15–24.e5 (2020).

38. Rix, G. et al. Scalable continuous evolution for the generation of diverse enzyme variants encompassing promiscuous activities. Nat. Commun. 11, 1–11 (2020).

39. Morrison, M. S., Podracky, C. J. & Liu, D. R. The developing toolkit of continuous directed evolution. Nat. Chem. Biol. 16, 610–619 (2020).

40. Molina, R. S. et al. In vivo hypermutation and continuous evolution. Nat. Rev. Methods Prim. 2, (2022).

41. Ravikumar, A., Arrieta, A. & Liu, C. C. An orthogonal DNA replication system in yeast. Nat. Chem. Biol. 10, 175–177 (2014).

42. Ravikumar, A., Arzumanyan, G. A., Obadi, M. K. A., Javanpour, A. A. & Liu, C. C. Scalable, Continuous Evolution of Genes at Mutation Rates above Genomic Error Thresholds. Cell. 175, 1946–1957.e13 (2018).

43. Zhong, Z., Ravikumar, A. & Liu, C. C. Tunable Expression Systems for Orthogonal DNA Replication. ACS Synth. Biol. 7, 2930–2934 (2018).

44. García-García, J. D. et al. Using continuous directed evolution to improve enzymes for plant applications. Plant Physiol. (2021) doi:10.1093/plphys/kiab500.

45. Jensen, E. D. et al. Integrating continuous hypermutation with high-throughput screening for optimization of cis,cis-muconic acid production in yeast. Microb. Biotechnol. 14, 2617–2626 (2021).

46. Javanpour, A. A. & Liu, C. C. Evolving Small-Molecule Biosensors with Improved Performance and Reprogrammed Ligand Preference Using OrthoRep. ACS Synth. Biol. 10, 2705–2714 (2021).

47. Wellner, A. et al. Rapid generation of potent antibodies by autonomous hypermutation in yeast. Nat. Chem. Biol. 17, 1057–1064 (2021).

48. Harvey, E. P. et al. An in silico method to assess antibody fragment polyreactivity. Nat. Commun. 13, (2022).

49. Vallina Estrada, E., Zhang, N., Wennerström, H., Danielsson, J. & Oliveberg, M. Diffusive intracellular interactions: On the role of protein net charge and functional adaptation. Curr. Opin. Struct. Biol. 81, (2023).

50. Brown, T., Hunter, W. N., Kneale, G. & Kennard, O. Molecular structure of the G·A base pair in DNA and its implications for the mechanism of transversion mutations. Proc. Natl. Acad. Sci. U. S. A. 83, 2402–2406 (1986).

51. Luria, S. E. & Delbrück, M. Mutations of Bacteria From Virus Sensitivity To Virus Resistance. Genetics 28, 491–511 (1943).

52. Foster, P. L. Methods for Determining Spontaneous Mutation Rates. Methods Enzymol. 409, 195–213 (2006).

53. Jain, M., Olsen, H. E., Paten, B. & Akeson, M. The Oxford Nanopore MinION: delivery of nanopore sequencing to the genomics community. Genome Biol. 17, 1–11 (2016).

54. Zurek, P. J., Knyphausen, P., Neufeld, K., Pushpanath, A. & Hollfelder, F. UMI-linked consensus sequencing enables phylogenetic analysis of directed evolution. Nat. Commun. 11, 1–10 (2020).

55. Karst, S. M. et al. High-accuracy long-read amplicon sequences using unique molecular identifiers with Nanopore or PacBio sequencing. Nat. Methods. 18, 165–169 (2021).

56. Lang, G. I. & Murray, A. W. Estimating the per-base-pair mutation rate in the yeast Saccharomyces cerevisiae. Genetics 178, 67–82 (2008).

57. Tian, R. et al. Engineered bacterial orthogonal DNA replication system for continuous evolution. Nat. Chem. Biol. (2023) doi:10.1038/s41589-023-01387-2.

58. Orr, H. A. The Rate of Adaptation in Asexuals. 968, 961–968 (2000).

59. Gerrish, P. J., Colato, A. & Sniegowski, P. D. Genomic mutation rates that neutralize adaptive evolution and natural selection. J. R. Soc. Interface 10, (2013).

60. Dunn, M. F. Allosteric regulation of substrate channeling and catalysis in the tryptophan synthase bienzyme complex. Arch. Biochem. Biophys. 519, 154–166 (2012).

61. Buller, A. R. et al. Directed evolution of the tryptophan synthase β-subunit for stand-alone function recapitulates allosteric activation. Proc. Natl. Acad. Sci. 112, 14599–14604 (2015).

62. Buller, A. R. et al. Directed Evolution Mimics Allosteric Activation by Stepwise Tuning of the Conformational Ensemble. J. Am. Chem. Soc. 140, 7256–7266 (2018).

63. Maria-Solano, M. A., Iglesias-Fernández, J. & Osuna, S. Deciphering the Allosterically Driven Conformational Ensemble in Tryptophan Synthase Evolution. J. Am. Chem. Soc. 141, 13049–13056 (2019).

64. Orij, R., Postmus, J., Beek, A. Ter, Brul, S. & Smits, G. J. In vivo measurement of cytosolic and mitochondrial pH using a pH-sensitive GFP derivative in Saccharomyces cerevisiae reveals a relation between intracellular pH and growth. Microbiology 155, 268–278 (2009).

65. Wennerström, H., Estrada, E. V., Danielsson, J. & Oliveberg, M. Colloidal stability of the living cell. Proc. Natl. Acad. Sci. U. S. A. 117, 10113–10121 (2020).

66. Xiang, L., Yan, R., Chen, K., Li, W. & Xu, K. Single-Molecule Displacement Mapping Unveils Sign-Asymmetric Protein Charge Effects on Intraorganellar Diffusion. Nano Lett. 23, 1711–1716 (2023).

67. Leander, M., Yuan, Y., Meger, A., Cui, Q. & Raman, S. Functional plasticity and evolutionary adaptation of allosteric regulation. Proc. Natl. Acad. Sci. U. S. A. 117, 25445–25454 (2020).

68. Shaw, A., Spinner, H., Shin, J., Gurev, S. & Rollins, N. Removing bias in sequence models of protein fitness. (2023).

69. Notin, P. et al. TranceptEVE: Combining Family-specific and Family-agnostic Models of Protein Sequences for Improved Fitness Prediction. bioRxiv 2022.12.07.519495 (2022).

70. Hie, B. L. et al. Efficient evolution of human antibodies from general protein language models. Nat. Biotechnol. (2023) doi:10.1038/s41587-023-01763-2.

