## Supplementary Materials for "Continuous evolution of user-defined genes at 1-million-times the genomic mutation rate"

### Materials and Methods

#### *DNA plasmid construction*

Plasmids used in this study are listed in Table S3, along with sources for DNA templates. Complete maps for these plasmids are made available for download on github at https://github.com/liusynevolab/OrthoRep_Rix_2023. All DNA templates for PCR were derived from previous studies or gBlocks (IDT). All primers were synthesized by IDT. All relevant primer pairs are listed in Table S4. Amplicons for construction of clonal plasmids were generated using Q5 Hot Start High-Fidelity DNA Polymerase (NEB). All non-library plasmids were constructed using Gibson Assembly and transformed into chemically competent *E. coli* strain TOP10 (ThermoFisher). Clonal plasmids were sequence verified by either Sanger sequencing (Azenta) or whole plasmid sequencing (Primordium).

#### *DNA library construction*

Amplicons for TP-DNAP1 libraries were generated using error prone PCR with GeneMorph II (Agilent) according to manufacturer instructions, aiming for ~3-5 nucleotide substitutions per sequence. Amplicons for all other libraries were generated with Q5 Hot Start High-Fidelity DNA Polymerase (NEB). For epPCR 1, the resulting PCR product was assembled into plasmids using Gibson assembly in 20 µL reaction volumes. For all other libraries, resulting PCR products were assembled into plasmids with Golden Gate assembly with T4 DNA ligase and BsaI-HF v2 or PaqCI (all NEB) and in a 40 µL reaction volume. Gibson reactions were run at 50 °C for 1 hour. Golden gate reactions were run isothermally at 37 °C for 1 hour and heat inactivated at 65 °C for 10 minutes. Reactions were purified with AMPure XP beads (Beckman), typically with a 0.9:1 bead:sample ratio according to manufacturer instructions. Libraries were transformed into high-competency electrocompetent *E. coli* TOP10 cells (ThermoFisher).

#### *Yeast strains, media, transformations, and DNA extraction*

All yeast strains used in this study and their provenance are listed in Table S5. Yeast were grown in liquid or on plates at 30 °C in synthetic complete (SC) growth medium (20 g/L dextrose, 6.7 g/L yeast nitrogen base w/ ammonium sulfate w/o amino acids (US Biological), appropriate nutrient drop-out mix (US Biological), as directed) or MSG SC growth medium (20 g/L dextrose, 1.72 g/L yeast nitrogen base w/o ammonium sulfate w/o amino acids (US Biological), appropriate nutrient drop-out mix (US Biological), as directed, 1 g/L L-Glutamic acid monosodium salt hydrate (ThermoFisher)) minus nutrients (referred to as -X where X is either the single letter amino acid code for an amino acid nutrient, or U for uracil) required for appropriate auxotrophy selection(s). Where selection for MET15 was required, cells were propagated in media lacking both methionine and cysteine. 500 µL liquid yeast cultures in 96-well deep well plates were incubated with shaking at 750 rpm. All other liquid yeast cultures were incubated with shaking at 200 rpm.

Yeast transformations, including p1 integrations and polymerase replacement integration, were performed as previously described[1]. For all integration transformations, plasmid DNA was linearized prior to transformation using either ScaI-HF or EcoRI-HF (both NEB) for p1 or genomic integrations, respectively. Due to its repetitive nature, deletion of FLO1 was performed by a URA3 knock-in knock-out method[2] (see Table S3, pFLO1-KO). Genetic deletions for TRP5 and MET15 were performed as previously described[3], using spacer sequences TTTGAGCCTGATCCCACTAG and GCTAAGAAGTATCTATCTAA, respectively. When isolating individual clones from genetic deletion and integration transformations, colonies were restreaked onto media agar plates of the same formulation to ensure isolation of only cells that have the desired genetic change.

All p1 plasmid sequences were generated by first generating a strain harboring a 'landing pad' p1 via integration and then integrating over this landing pad to generate the desired p1 construct. To enable construction of the landing pad strain, the wt TP-DNAP1 was integrated at the CAN1 locus using pGR475. A sequence encoding a partial LEU2 sequence lacking the N terminus was then integrated over the wt p1 using pGR420 to generate the landing pad p1. The wt p1, which encodes the TP-DNAP1, was then cured out via 3-4 1:1000 passages. The p1 plasmid(s) encoding the desired sequence were then generated via integration using cassette(s) that include a LEU2 sequence lacking the C terminus (*e.g.* pGR438). The overlap between the LEU2 on the landing pad and the new integration cassette was used to reconstitute full length LEU2 only when integration occured on p1, reducing the likelihood of genomic integration.

The polymerase replacement integration transformation was performed by first digesting 0.5-2 µg of the polymerase replacement plasmid or library with EcoRI-HF in a 25 µL reaction per 1x transformation followed by directly transforming this digestion reaction into a yeast strain encoding the CAN1-WT-TP-DNAP1 landing pad (all polymerase libraries were transformed into OR-Y488). Library scale transformations were carried out at 20-40x scale. Transformed yeast were plated onto solid MSG SC -LR or -MCR media w/ 100 mg/L nourseothricin (for positive selection of integration) and 200 mg/L L-canavanine (a toxic L-arginine analog for counterselection of cells that fail to perform polymerase replacement and remove the arginine permease CAN1). Leu or Met/Cys dropout was used to maintain selection for p1-encoded LEU2 or MET15, respectively, while Arg dropout was usedto improve L-canavanine selection.

Extraction of genomic DNA (gDNA) and p1/p2 plasmids was performed as previously described for 1.5 mL yeast culture volumes[1]. This procedure was used for all experiments except for DNA extracted for use in HTS dataset 6, which was instead performed in 96-well format for higher throughput. In brief, a 96-well block of 500 µL of saturated yeast cultures was centrifuged (2500 × G, 5 min), supernatant was discarded, pellets were resuspended in 1 mL 0.9% NaCl, this resupension was again centrifuged (2500 × G, 5 min), and the supernatant was discarded. The resulting pellet was resuspended in 250 µL Zymolyase solution (0.9 M D-Sorbitol (Sigma Aldrich), 0.1 M Ethylenediaminetetraacetic acid (EDTA, Sigma Aldrich), 10 U/mL Zymolyase (US Biological)) and incubated with shaking (37 °C, 200 RPM). The 96-well block was then centrifuged (2500 × G, 5 min), supernatant was discarded, and pellets were resuspended in 280.5 µL proteinase K solution (250 µL TE (50 mM Tris-HCl (pH 7.5), 20mM EDTA), 25 µL 10% sodium dodecyl sulfate (SDS, Sigma Aldrich), 5.5 µL proteinase K stock solution (10 mg/mL proteinase K (ThermoFisher)). The 96-well block was then incubated at 65 °C for 30 min, combined with 75 µL 5M potassium acetate (ThermoFisher), and incubated on ice for 30 min. The 96-well block was centrifuged at 12,000 × g for 10 min, the resulting supernatant was combined and mixed with 2 volumes buffer PB (5 M Guanidine hydrochloride (ThermoFisher), 30% isopropanol, 70% water), and this mixture was applied to a 96 well DNA-binding plate

(Epoch Life Science) on a vacuum manifold. Flow through was discarded, columns were washed with PE buffer (10 mM Tris-HCl (ThermoFisher), 80% ethanol, 20% water, pH 7.5), centrifuged and dried, and 60 μL water was applied to columns for elution by centrifugation (2500 × G, 5 min).

*High throughput sequencing*

All high throughput sequencing datasets are listed in Table S6, along with the method used to construct them. All PCRs for high throughput sequencing were performed with Platinum SuperFi II DNA Polymerase (ThermoFisher). For short read paired end sequencing, both low cost/yield (AmpliconEZ, Azenta) and high cost/yield (HiSeq paired end 150, Novogene) were performed directly on PCR products generated in either one or two rounds of PCR using primers that each included an adapter sequence, a 6- or 7-nucleotide barcode, or both.

For in-house long read sequencing, we used the Oxford Nanopore Technologies nanopore sequencing platform. Due to the lower accuracy of nanopore sequencing, we employed modified versions of previously described methods for the construction of DNA libraries that yield multiple reads of the same original DNA molecule, allowing for computational reconstruction of high accuracy sequences[4,5]. We refer to the first of these as *in vivo* downsampled unique molecular identifier (UMI) PCR, which was used for most nanopore sequencing in this study (Table S6). This involved first a 2-cycle “UMI tagging” reaction, in which primers (*e.g.*, primer pair 2) were used to append both UMIs and universal DNA sequences (for further amplification) to both ends of the target sequence with the following components:

- ~1-50 ng of purified yeast or *E. coli* miniprep
- 5 μL 2x SuperFi II master mix
- 1 μM each primer
- water to 10 μL

and with these thermocycler conditions:

1. 98 °C for 30 sec
2. 98 °C for 10 sec
3. 65 °C for 1 sec
4. 60 °C for 45 sec w/ ramp down from 65 to 60 @ 0.2 °C per second (this should result in 25 seconds of ramp down time and 20 seconds of hold time)
5. 72 °C extension, 1 min / kb
6. Go to step 2 (1x)
7. 72 °C for 2 min

Next, UMI primers were removed using ExoSAP-IT (ThermoFisher) according to manufacturer instructions. 10 μL of the resulting reaction was then used as template in a 25 μL Platinum SuperFi II PCR with 1 mM $MgCl_2$ supplemented, using primers (*e.g.*, primer pair 3) that included BsaI or PaqCI sites and 7 nucleotide barcodes in forward/reverse combinations that were unique for each sample. The resulting uniquely barcoded PCR products were then combined, purified using AMPure XP beads, used in a library Golden Gate reaction with an *E. coli* vector, then transformed into high competency *E. coli* as described above. Plasmid pGR554 (Table S3) was designed for this purpose and contains both CcdB and sfGFP, which both are replaced with the insert during Golden Gate assembly, as well as NotI and SbfI sites, strategically placed to enable separation of the desired library insert from the backbone prior to sequencing. CcdB and sfGFP provided counterselection and visualization of colonies resulting from undigested vector. (For reference, cloning into any *E. coli* vector suffices, so long as unique library members are associated with a unique relatively short (20-50 bp) sequence.) Resulting colonies each contained many copies of a unique plasmid species encoding a UMI-tagged library member. To obtain good coverage of each sequence and UMI with nanopore sequencing, the resulting library was downsampled by only harvesting ~20-fold fewer colonies than the expected number of reads, amounting to 100 – 200 thousand colonies for a standard MinION flow cell. Plasmid DNA from this library was then miniprepped, digested (for example with NotI-HF or NotI-HF/SbfI-HF (both NEB) if using plasmid pGR554), and gel extracted prior to sequencing.

Construction of the library for HTS dataset 8a was performed using a similar approach, but with a yeast expression vector, and with UMIs and library members inserted into the vector in two distinct steps so that UMIs are present at a single location in the plasmid and would not need to be immediately adjacent to library members, mitigating any potential effects of UMIs on expression. In brief, libraries of UMIs generated via PCR using primer pair 8 were cloned into plasmid pUMI by Golden Gate assembly with BsmBI-v2 and *E. coli* electrotransformation to generate an intermediate UMI library. Different variants of pUMIs with unique, known 7-nucleotide barcodes were used for each library to allow multiplexing. These libraries were then used as the vector into which evolved TrpB library members, PCR amplified using primer pair 9, were cloned via standard restriction cloning, with inserts digested with PaqCI and vectors digested with BsaI-HFv2. Isothermal ligation (1 hour 37 °C, 20 mins 65 °C) was then performed with T4 ligase in 40 µL reaction volumes, AMPure bead purified, and transformed into high efficiency *E. coli*. This was carried out for 6 unique libraries: downsampled in yeast with selection, downsampled in yeast without selection, and not downsampled in yeast, each for both timepoints (generation 350 and generation 540) chosen from TrpB evolution. The intermediate UMI libraries were generated at a library size of >100-fold larger than the desired final library size to minimize the chance that distinct library members would be tagged with identical UMIs.

The second method used for generating libraries for high accuracy nanopore sequencing was adapted from methods described in Volden *et al.*[5], Oliynyk and Church[6], and Zhang and Tanner[7]. It involved circularization of the target sequence and use of this circularized product as template for rolling circle amplification (RCA) using strand displacing DNA polymerases (Fig S15). In brief, UMI-tagged PCR products were generated as described above, albeit with complementary Type IIS cut sites (BsaI or PaqCI) on the forward and reverse primers used during PCR amplification such that the two ends of the amplicon ligated to each other during a Golden Gate assembly reaction, forming a circular product. Due to the large amount of template DNA required for rolling circle amplification, a large amount of amplicon was used in the circularization reaction, typically 1-2 pmols in 100 µL Golden Gate assembly reactions. Oliynyk and Church[6] provide a useful discussion of relevant considerations for such circularization reactions. An isothermal Golden Gate assembly reaction was then performed to circularize the amplicon library.

Following Golden Gate assembly, uncircularized DNA was digested with lambda exonuclease (NEB), exonuclease I (NEB), and exonuclease III (NEB) in a 1:1:0.1 ratio, which was added directly to the Golden Gate reaction in a 1:10 exonuclease mixture to sample ratio. This reaction was incubated at 37 °C for 45 min, then 80 °C for 15 min. The reaction was then AMPure bead purified with a 0.7:1 bead:sample ratio, eluting in a maximum of 10 µL of water, and the resulting circularized library was used in a rolling circle amplification reaction with the following components combined on ice:

- 4 µL 10X NEB buffer 4 (NEB)
- 4.8 µL 10 mM dNTPs
- 2.64 µL 5 U/µL Bsu DNA Polymerase, Large Fragment (NEB)
- 1.6 µL 10 µg/µLT4 gene 32 protein (NEB)
- 0.2 – 1 µg circularized DNA library
- RCA primers, 2 µM each (must bind internal to first primer set, *e.g.* primer pair 7)
- Water to 40 µL

This reaction was incubated at 37 °C for 3 hours. SDS-containing loading dye was added directly to the reaction, and the entire sample was run on an agarose gel. Bands corresponding to 3x–6x concatemers were gel extracted, and purified DNA was used for nanopore sequencing. Unlike RCA using Phi29, this method enabled size selection of specific repeat numbers and did not require a 'debranching' step, but required large amounts of input DNA and in our experience suffered from high sensitivity to DNA contamination. Use of Phi29 with random hexamers, followed by debranching, is therefore a reasonable alternative.

Following construction of *in vivo* downsampled UMI PCR or RCA libraries, nanopore library preparation and sequencing was performed using the most up-to-date ligation sequencing kit (*e.g.* LSK-114) and flow cell (*e.g.* R10.4.1), following manufacturer instructions, with two exceptions. First, ½ volumes (but unaltered DNA input) for end prep and ligation reactions were used and second, FFPE Repair Mix was not used during end prep reactions.

All relevant high throughput sequencing datasets will be made available on the NCBI sequence read archive (SRA) prior to peer-reviewed publication.

*Error rate measurement by mutation accumulation*

Following a polymerase replacement integration transformation, colonies were picked into liquid media of the same media formulation as that used for selection after transformation, and then grown to saturation. The resulting saturated culture was miniprepped to serve as the $0^{th}$ passage, $p_0$. The culture was then also propagated in the appropriate growth medium for maintenance of the orthogonal plasmid, typically SC -L, using a dilution factor $d$ (typically 128, 256, or 512) that is consistent throughout the experiment. At least one additional saturated culture from this time course was miniprepped to serve as the $t^{th}$ passage, $p_t$. We used only two passages/timepoints in all mutation accumulation error rate measurement experiments except that which produced HTS dataset 6. The number of generations, $g$, that separated $p_0$ from $p_t$ was used to calculate the mutation rate and can be approximated as

$$g_t = t \times \log_2 d\,.$$

We note that this approximation assumes no cell death and equivalent saturation at each passage. Cultures were manually propagated several times until a total number of generations of at least 50 was reached. Miniprepped yeast DNA for each timepoint was used as template

DNA for high throughput sequencing, and custom scripts, organized within the Maple pipeline, were used to calculate mutation rates.

Mutation rates were calculated as the rate of accumulation of a mutation type (all substitutions, individual substitutions, insertions, or deletions), $j$. We denote $\mu_j$ as this rate calculated for mutation type $j$. High accuracy HTS was used to first obtain $c_{j,t}$, the total counts of mutation $j$ (*e.g.*, A to T) among all sequences in passage $t$. For substitution mutations, to account for the influence of variable A/T/G/C content in the sequence being analyzed, this count is normalized to obtain $n_{j,t}$, the expected count for an idealized sequence with a 1:1:1:1 A:T:G:C ratio. For a substitution at a particular nucleotide where the nucleotide occurs $w$ times within a reference sequence of length $L$, $n_{j,t}$ is calculated as

$$n_{j,t} = \frac{c_{j,t}}{w} \times \frac{L}{4}\,.$$

The total normalized count of all substitution mutations at each timepoint is then calculated as the sum of all $n_{j,t}$ for all twelve substitution types. However, for insertions and deletions, $n_{j,t}$ is not normalized in this way, and is instead equivalent to $c_{j,t}$, the total number of nucleotides inserted or deleted among all sequences for that timepoint.

To obtain the per-nucleotide, per-generation mutation rate $\mu_j$, we used the total number of sequences analyzed for each timepoint, $s_t$, to calculate the per-nucleotide frequency of mutation $j$ for each timepoint. We then use linear regression on these normalized per-nucleotide frequencies according to

$$\frac{n_{j,t}}{L \times s_t} = \mu_j \times g_t + b_j,$$

where $\mu_j$ and $b_j$ are the slope and intercept, respectively, of the best fit line for all $t$ timepoints. When the number of generations between the 0$^{th}$ timepoint and the initiation of mutagenesis (typically polymerase replacement) is accurately estimated and no mutations fully fixed within the population prior to the first timepoint, $b_j$ should be close to 0. Regardless, we do not report $b_j$, as it has no bearing on $\mu_j$ across experiments. We report $\mu_j$ as the per-base per-generation rate of accumulation of mutation type $j$. Mutation tabulation was performed by the script mutation_analysis.py and all other operations related to mutation rate calculation were performed by the script plot_mutation_rates.py, both of which are contained within the Maple pipeline. All reported mutation rates were calculated using mutation tabulation within a sequence region that was not under functional selection.

*TP-DNAP1 library selection and screening*

Following TP-DNAP1 replacement library construction, resulting purified plasmid DNA was used for a polymerase replacement integration transformation. Prior to transformation, OR-Y488 was grown up in SC -L + 1 mg/L 5-fluoroorotic acid (US Biological) for counterselection against cells that had reverted the inactivating mutation in ura3* by chance. Following library transformation,plating on media selecting for cells that had replaced the wild type TP-DNAP1 with library variants, and 48 hour incubation, colonies were harvested in bulk and immediately plated onto SC -LU plates. These plates were then incubated for 48 hours, and resulting colonies were either picked into liquid media for mutation rate or frequency characterization (by mutation accumulation or fluctuation assays, respectively) or were harvested in bulk and immediately plated onto SC -LUW media. After colony formation, colonies were either picked into liquid media for mutation rate characterization by mutation accumulation or were harvested in bulk, miniprepped for gDNA isolation, and used as template for a non-mutagenic PCR to generate amplicons for Golden Gate assembly into pGR554, which was then retransformed into OR-Y488 to repeat the selection and perform mutation rate screening.

The fluctuation test was performed as follows. Following transformation of the epPCR 1 TP-DNAP1 library into OR-Y488 and selection for ura3* reversion on solid media, individual colonies were picked from this plate, inoculated into 500 µL SC -LU media in a 96 well block, and grown to saturation. Cultures were then passaged 1:10,000 into 200 µL SC -LU, 12 replicates per each individual colony, and grown to saturation. Cultures were centrifuged, washed with 0.9% NaCl, and pellets were resuspended in 35 µL 0.9% NaCl. Each replicate 10 µL of this resuspension was plated onto SC -LUW plates. A subset of cultures were titered and plated on SC -LU plates to estimate population size. Plated cells were allowed to grow for 4 days, and revertants on each spot were counted. Counts were used to estimate the *m* value using the FALCOR online web tool (https://lianglab.brocku.ca/FALCOR/) and the Ma-Sandri-Sarkar Maximum Likelihood Estimator. Mutation frequency was calculated from this *m* value as previously described[8], using a target size of 1 (only one mutation is capable of restoring Trp5 activity). Copy number was not considered and therefore per base substitution rate was not calculated using this method.

*TrpB evolution*

Plasmid pGR595 (TP-DNAP1 BadBoy2) was first transformed into yeast strain OR-Y484 according to the polymerase replacement procedure described above. The resulting strain (OR-Y538) was then transformed with plasmid pGR438 (TrpB with lineage barcodes), following the p1 integration procedure described above, plating on SC -L media. All ~400 resulting colonies were harvested together and passaged into 512 µL SC -L media in all wells of a 96-well block and grown to saturation. DNA extracted from these resulting cultures served as passage/timepoint 0, which we approximate to be ~50 generations from TrpB p1 integration. These cultures were also passaged 1:1024 (0.5 µL into 512 µL) for all passages in the experiment into growth media and with timepoints taken as described in Fig. S14. DNA extraction for timepoints was performed by combining all 96 saturated cultures for a specific timepoint and extracting DNA from the pooled cultures according to the 1.5 mL DNA extraction protocol.

To downsample evolved populations for cloning UMI-tagged TrpB libraries, yeast cultures from the passages corresponding to generation 340 and 510 were inoculated into SC -L media from glycerol stock and grown to saturation. Saturated cultures were then combined, and multiple serial dilutions of both cultures were plated onto SC -L media. Plates derived from generation 340 and 510 with ~3700 and ~1700 colonies respectively were harvested. These cultures were passaged 1:1000 into 2 mL of either SC -LW + 400 µM indole (Sigma-Aldrich) (selective) or SC -L (nonselective) growth media and allowed to grow to saturation. This process was repeated twice more for each of the two media. All three of the resulting saturated cultures for each selective and nonselective conditions were pooled and DNA was extracted to serve as template for cloning the TrpB library fitness assay library, in addition to DNA extracted from generation 340 and 510 without downsampling. Note that this downsampling performed in yeast preceded the downsampling in *E. coli* necessary for proper sequencing coverage.

*Pooled TrpB fitness assay*

UMI-tagged evolved TrpB libraries and equivalent plasmids expressing two control TrpB sequences (*Tm*Triple and TrpB-003-1-A) were transformed into yeast strain OR-Y260 and plated onto -LH media, resulting in 40-fold coverage of the ~120 thousand member library. Colonies were harvested and the library was spiked with each of the two control TrpB-expressing yeast strains at a 1:1000 control:library ratio. DNA was extracted from the resulting library to serve as the 0$^{th}$ timepoint. This library was also passaged 1:100 into 50 mL of either SC -L (nonselective), SC -LW + 400 µM indole (weakly selective), or SC -LW + 25 µM indole (strongly selective) and grown to saturation. This passaging and growth was repeated for each of the three growth conditions five times for a total of six passages. Of these, DNA was extracted from passages 1, 2, 5, and 6 to serve as additional timepoints. DNA from all timepoints were used as templates for PCR amplification of only the UMI and barcode regions for high throughput sequencing.

Enrichment scores were calculated following the procedure described in Rubin *et al.*[9], albeit with two modifications to account for disparate sequencing coverage over multiple timepoints. First, counts of each UMI at each timepoint were normalized to the average count of all UMIs that persisted throughout all timepoints prior to log transformation and weighted linear regression. Second, regression weights were multiplied by this average count. All enrichment calculations were performed by the Maple pipeline within the script enrichment.py.

*High-throughput sequencing analysis*

High accuracy consensus sequence generation, alignment, demultiplexing, genotype identification, mutation rate analysis, and basic dataset visualization were performed by version v0.10.4 of the Maple pipeline, which uses the Snakemake workflow management library, and is available on https://github.com/gordonrix/maple. Parameters and settings for Maple analyses for each dataset and all other code used for analysis are available on Github (https://github.com/liusynevolab/OrthoRep_Rix_2023).

*TrpB fitness prediction with TranceptEVE*[10]

A multiple sequence alignment (MSA) of natural TrpB subunits was created using 5 iterations of jackhmmer[11] to query the UniRef100 database with a bitscore of 0.9. Columns with more than 20% gaps were ignored, and we used a theta parameter of 0.8 to downweight sequences with more than 80% sequence homology, as described in Hopf *et al.*[12]. We trained 4 separate EVE models using this MSA and used them to calculate the log probabilities of the mutated sequences. These scores were ensembled with the Tranception log probabilities for each sequence by taking a weighted average of 60% Tranception score and 40% EVE score.

*Null hypothesis sequence simulation and analysis*

A dataset of TrpB variants that contained random mutations representative of biases due to the relevant mutation preferences and wild type sequence, but that were not subject to selective forces, was generated through a simple simulation. First, the number of synonymous mutations within the ORF of each unique evolved TrpB sequence was approximated as the number of nucleotide mutations minus the number of nonsynonymous mutations. For each real unique genotype, a corresponding simulated genotype was generated by starting from the wild type TrpB sequence used in the evolution experiment and stochastically sampling nucleotide mutations with probabilities determined by the mutation rates of the same polymerase used for TrpB evolution (BadBoy2) until the same number of synonymous mutations as the evolved genotype was reached. Additional information such as the timepoint from which the real sequence was identified and the count of the genotype was also replicated for the corresponding simulated sequence. To account for some minor strand-dependent mutational biases, mutation rates and spectrum calculated for a sequence in the same position and orientation (relative to the LEU2 gene) as TrpB were used to generate the simulated sequences.

Simulated and evolved sequences were analyzed identically. Isoelectric point and hydrophobicity index (gravy) were both calculated using the ProtParam module in BioPython[13]. Mesophilic adaptation mutations were selected from Haney *et al*.[14] as the inverse of the 17 mesophile to thermophile “Replacements most biased in number” with $P<0.005$. Alternative sets of amino acid replacements were not evaluated.

*Computational lineage downsampling*

Lineage barcodes were identified using the demultiplexing feature within the Maple pipeline. The 100 most frequently observed lineage barcodes only from the first timepoint of TrpB evolution were used for all lineage analyses. Lineage barcodes for all remaining timepoints were assigned from this list of 100 barcodes, allowing for a nucleotide hamming distance of 1. For global covariation analysis, all barcodes appearing in at least 100 sequences were identified, and 100 sequences from each lineage were randomly extracted for analysis of mutation frequencies. Covariation with specific mutations was performed similarly, except that only sequences that contained the specific mutation and were identified from specified timepoints were considered prior to randomly extracting sequences for analysis of mutation frequencies. To ensure random sampling did not bias results, this process was performed multiple times with virtually identical results.

### Supplementary Figures

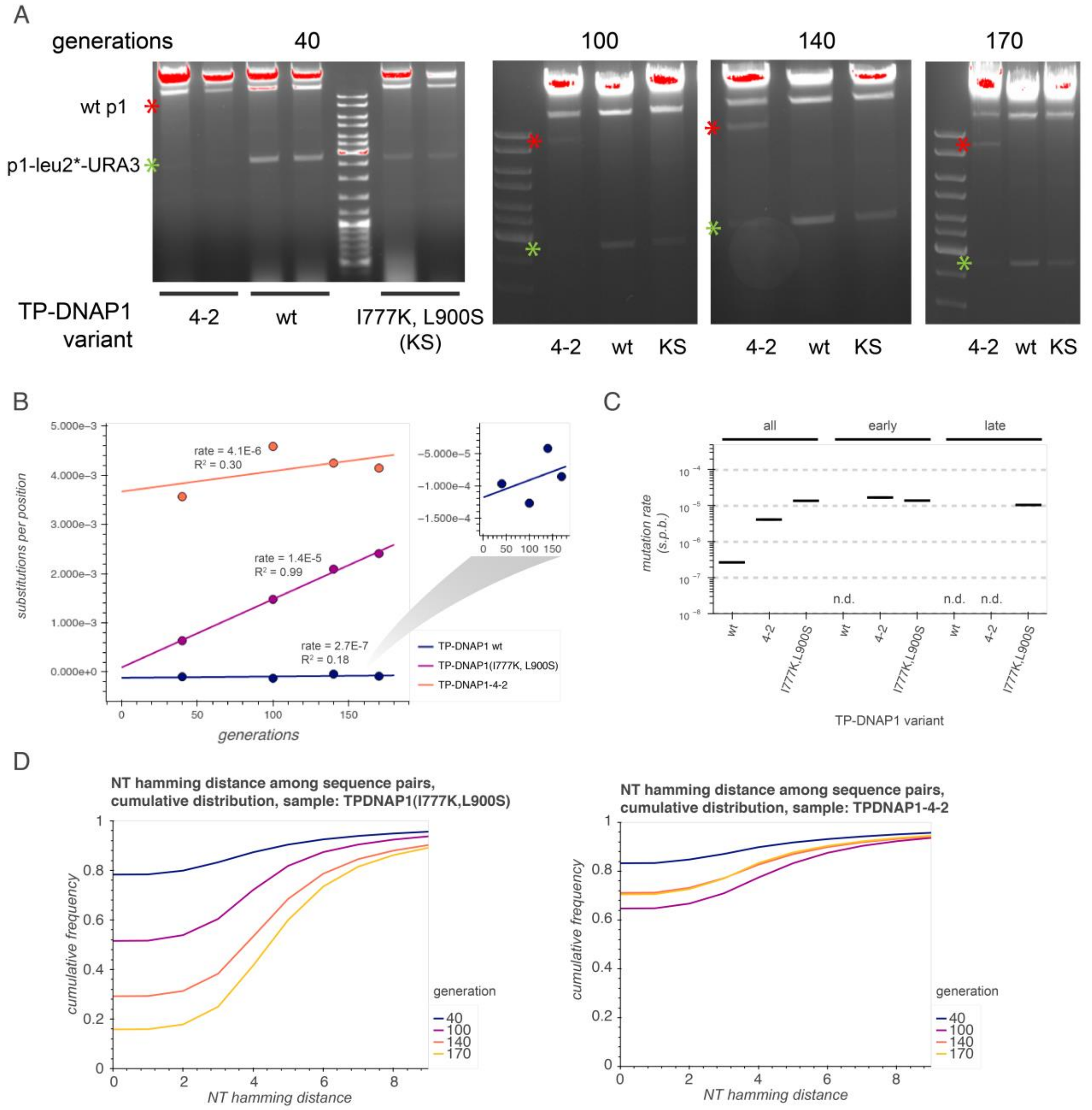


**Fig. S1. Validation of mutation accumulation and comparison of p1 maintenance by legacy TP-DNAP1 variants.** (**A-D**) Yeast strains encoding p1-leu2*-URA3 were transformed with plasmids encoding wild-type (wt) TP-DNAP1, TP-DNAP1-4-2, or TP-DNAP1(I777K, L900S) and passaged for 130 generations under selection for URA3 to allow for accumulation of mutations in leu2*. DNA was isolated from these samples at four timepoints throughout the experiment. Gel electrophoresis of these DNA samples (A) revealed the resurgence of a DNA band corresponding in length to wt p1 (~9 kb) in the TP-DNAP1-4-2 sample, but not others.

Timepoint DNA isolates were used for PCR amplification, high throughput sequencing, and mutation analysis of a ~450 bp region demonstrated that TP-DNAP1(I777K, L900S) maintained both a consistent mutation rate (B, C) and monotonic diversification throughout the experiment (D). Two replicates were isolated for the first timepoint for gel electrophoresis, but only one of these was isolated for subsequent timepoints and used for mutation accumulation. Mutations per base for WT is shown rescaled in (B) to highlight the poor linear fit. n.d., not detected.

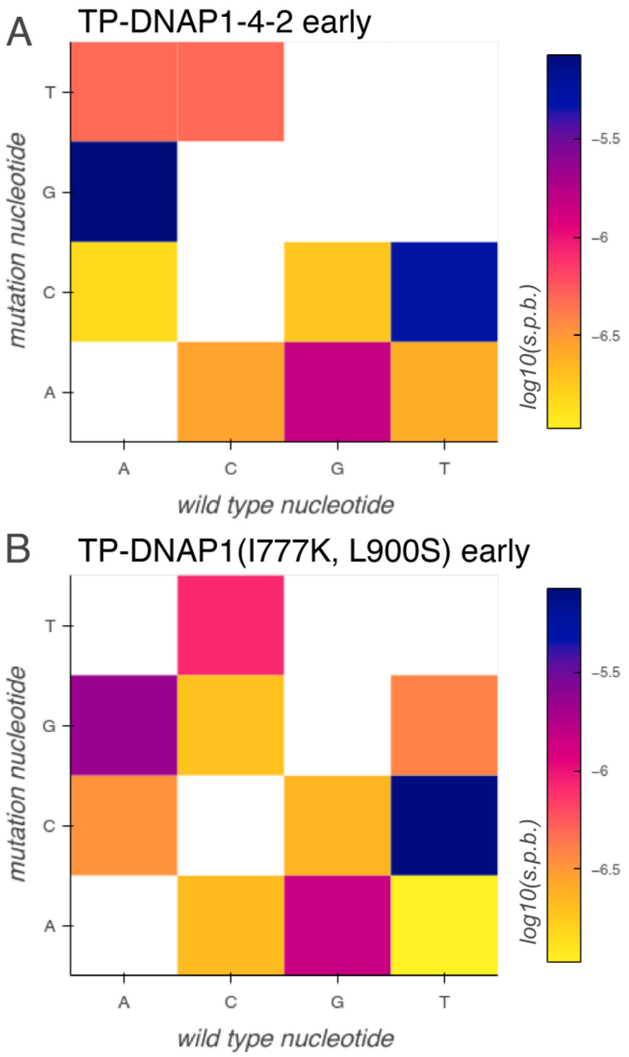


**Fig. S2. Mutation spectrum of legacy TP-DNAP1s.** (**A-B**) Heatmaps of log transformed individual substitution rates for all 12 substitution types for TP-DNAP1-4-2 (A) and TP-DNAP1(I777K, L900S) (B) from the first two timepoints of mutation accumulation in HTS dataset I (~60 generations, see Fig. S1).

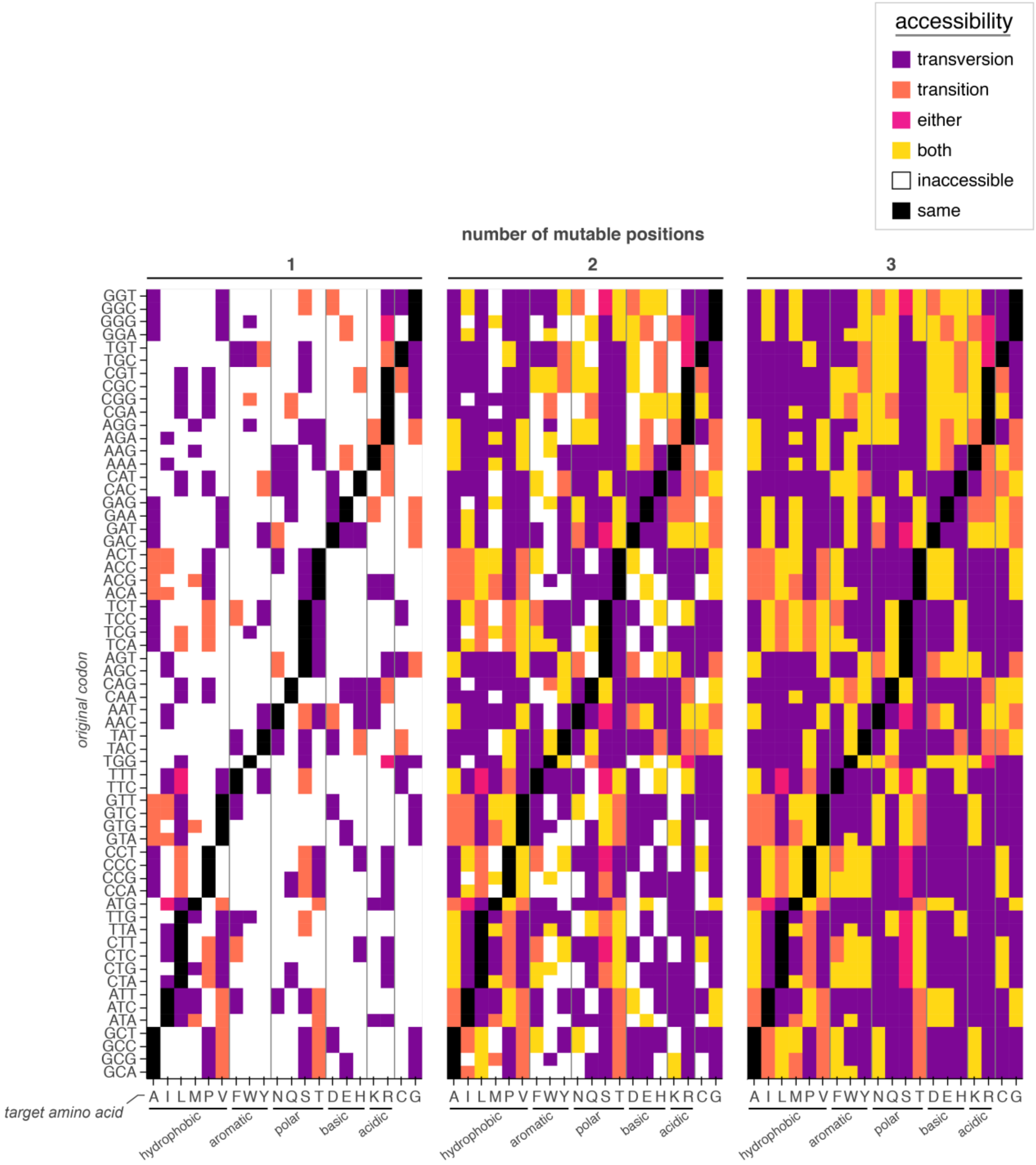


**Fig. S3. Amino acid accessibility by mutation type.** Accessibility of any codon for each amino acid from each of the 64 possible codons by one (left column), two (middle column), or three (right column) nucleotide mutations. Color indicates whether the codon can reach the corresponding amino acid with only transversions (purple), with only transitions (salmon), either with only transversions or with only transitions (pink), only with both transversions and transitions (yellow), or is inaccessible (white) with at most the indicated number of mutable positions. Multiple mutations to the same position within the codon were not considered.

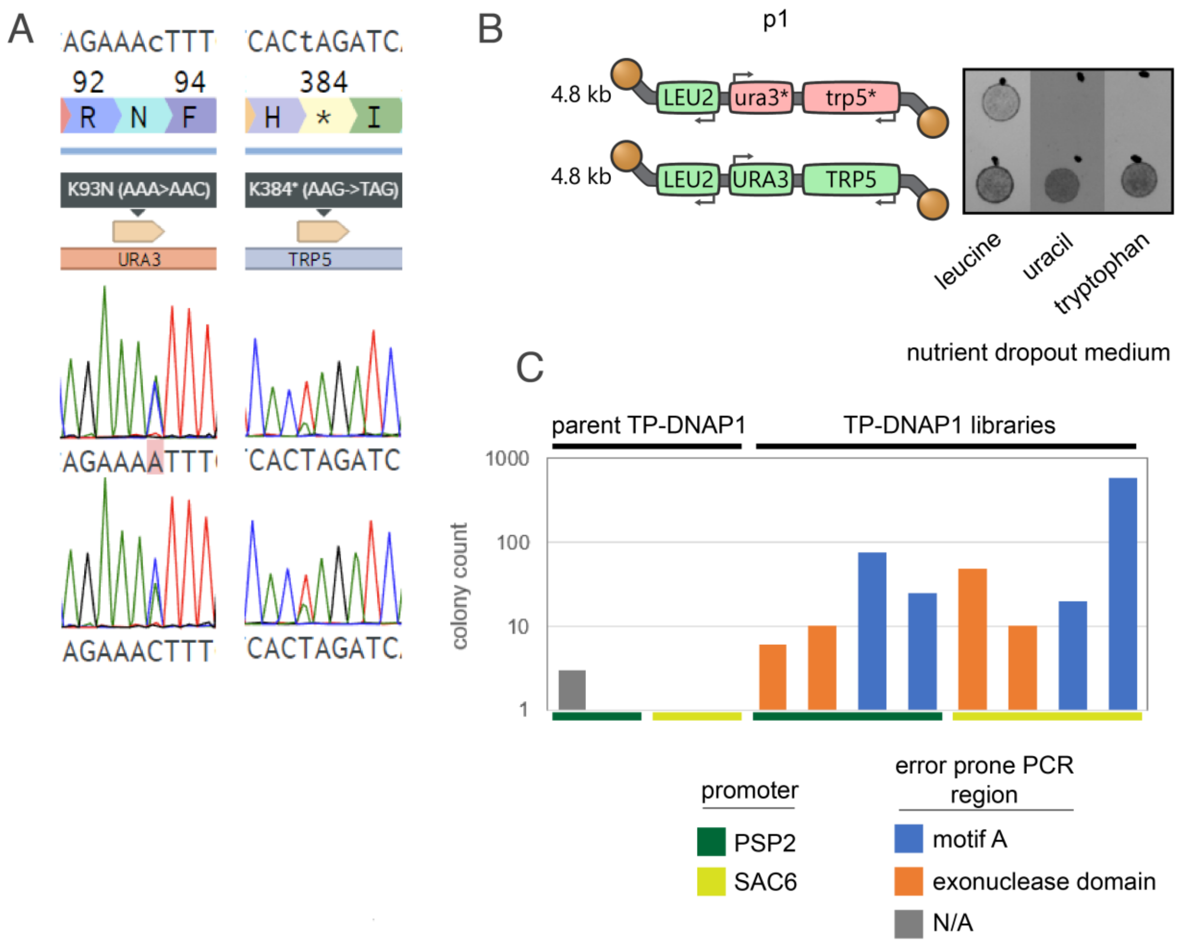


**Fig. S4. Validation of transversion-specific selection and epPCR 1.** (**A**) Sanger sequencing of the ura3-K93N (ura3*) or trp5-K384* (trp5*) transversion mutations in two distinct clonal populations of yeast harboring the p1-ura3*-trp5* plasmid following selection in -uracil or -tryptophan media, respectively. In all cases, the only visible peaks were those corresponding to the original sequence or the desired transversion mutation. 100% fixation of the expected mutation was not enforced due to the multicopy nature of p1. (**B**) Plating assay demonstrating that p1-encoded auxotrophy markers URA3 and TRP5 were rendered inactive by the K93N and K384* mutations to catalytic lysines. (**C**) URA3 revertant colony counts for either TP-DNAP1 epPCR 1 libraries or the parent TP-DNAP1(I777K, L900S) transformed into OR-Y488. TP-DNAP1(I777K, L900S) was used as the template sequence for error prone PCR mutagenesis of one of two regions of the polymerase expressed under one of two promoters for a total of four $10^3$ – $10^4$ member TP-DNAP1 libraries which were then transformed into the OR-Y488 strainharboring p1-ura3*-trp5*. Two biological replicates of each of these libraries were subject to selection for URA3 alongside mock libraries encoding only the parental TP-DNAP1 sequence and resulting revertants were quantified by titering and replating.

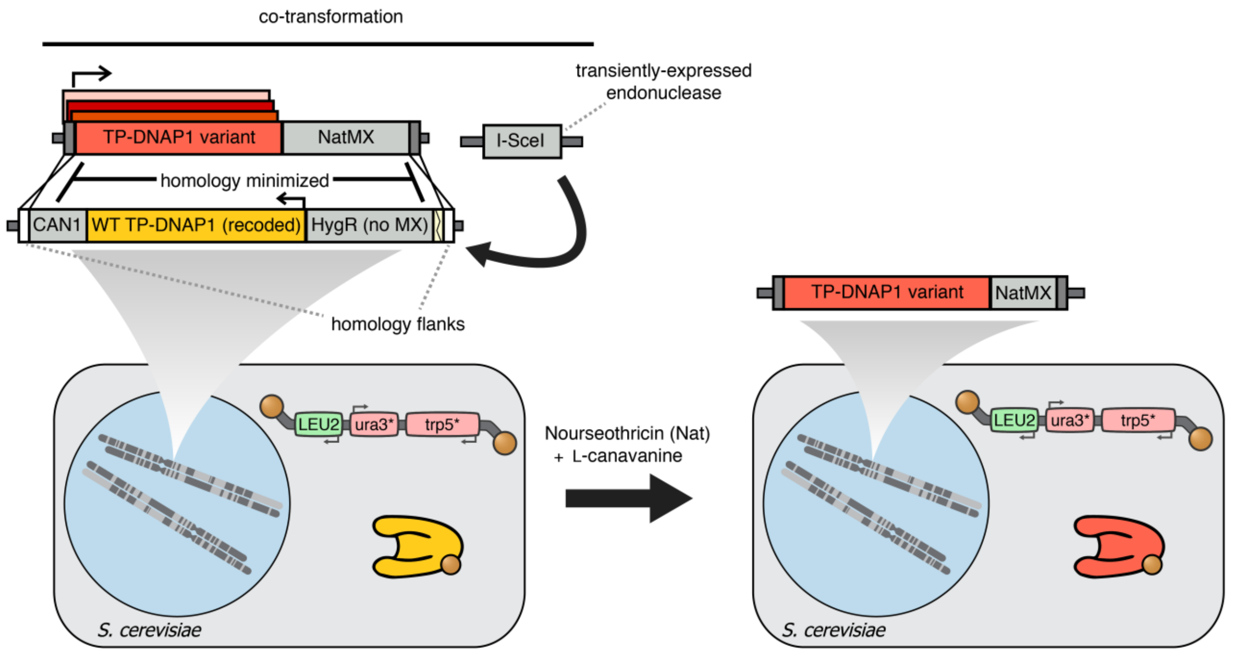


**Fig. S5. Illustration of the design of the polymerase replacement transformation.** A strain encoding a "landing pad" DNA sequence that includes the wild type (WT) TP-DNAP1 and an I-SceI cut site at the CAN1 locus was co-transformed with both a TP-DNAP1 variant (typically in library format) and a transient I-SceI endonuclease expression cassette[15]. Integration of NatMX was selected for using nourseothricin. Retention of the CAN1 in the landing pad was selected against using L-canavanine.

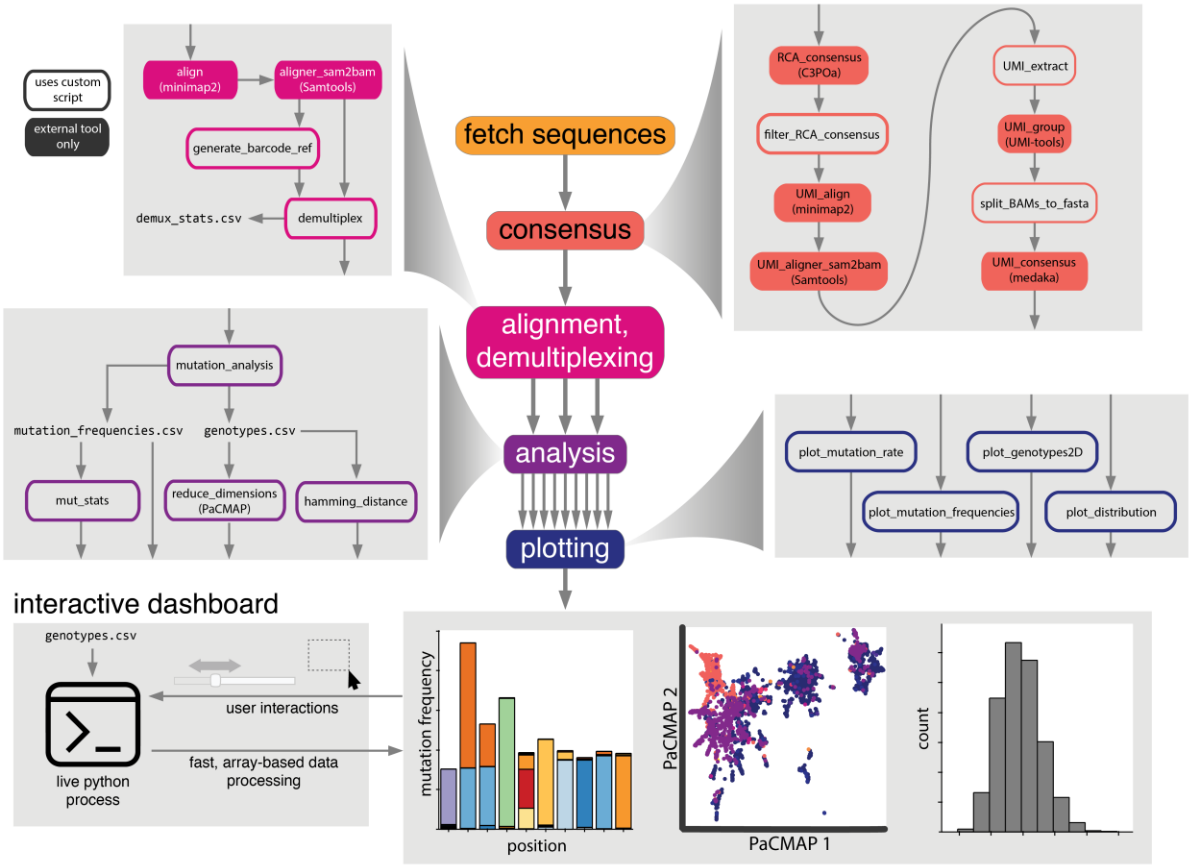


**Fig. S6. Flowchart of the programmatic steps performed by the mutation analysis for parallel laboratory evolution (Maple) pipeline.** To facilitate rapid exploration and analysis of high throughput sequencing datasets, we built an end-to-end data processing pipeline that takes as input a sequencing dataset and a minimal set of additional user inputs and carries out the necessary steps to produce commonly desired visualizations as well as the data that supports those visualizations. This includes generating high accuracy consensus sequences from multiple reads of a sequence via rolling circle amplification (RCA) or unique molecular identifiers (UMI), alignment-based demultiplexing to separate and label sequences derived from different samples, and mutation analysis to generate human-readable .csv outputs that are further analyzed and visualized by Maple or can be viewed and analyzed by the user. Maple also includes an interactive dashboard that allows for user interaction with visualizations, such as selection of sequences that cluster together when plotted by the output of the dimensionality reduction tool PaCMAP.

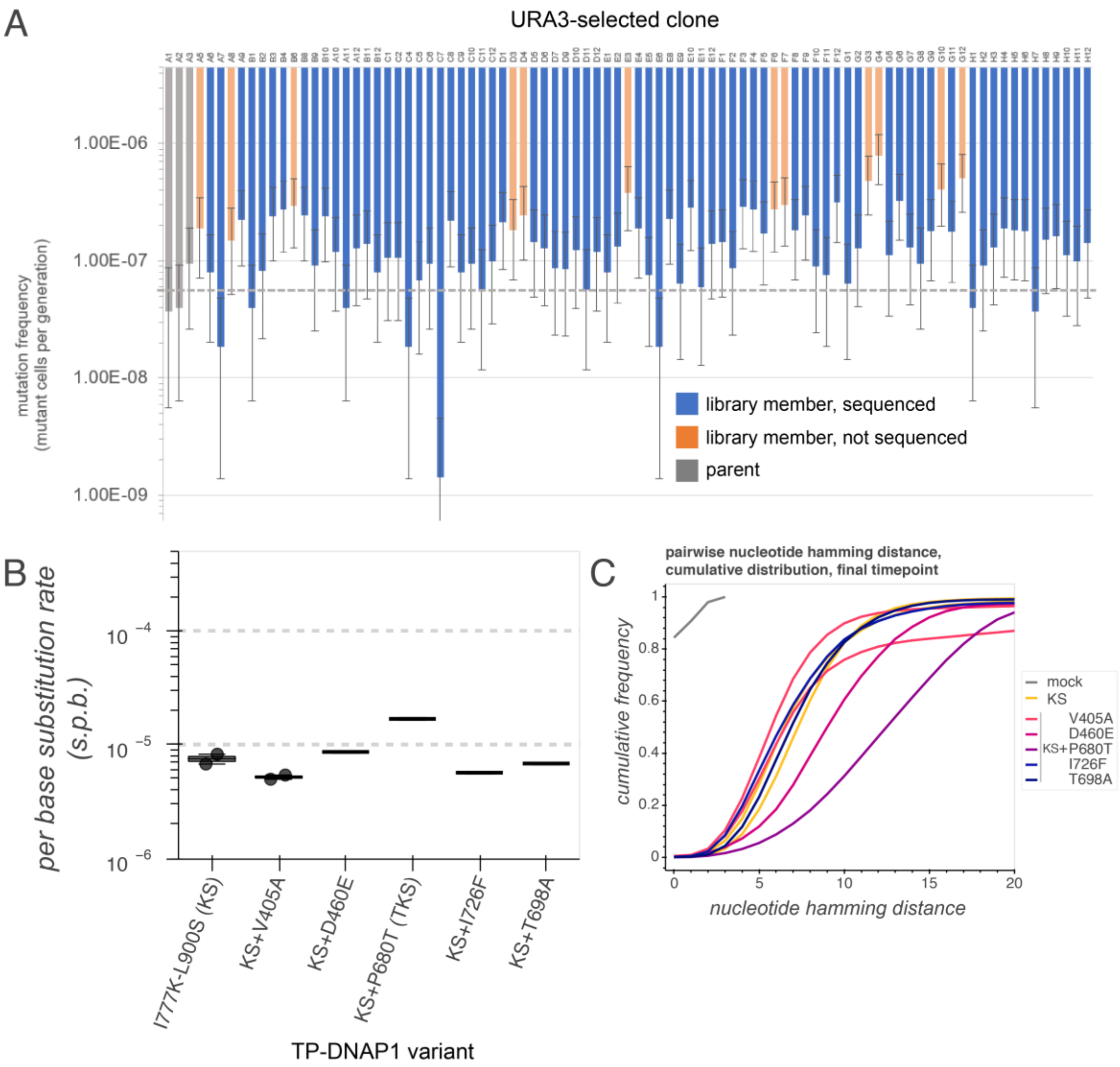


**Fig. S7. Identification of TP-DNAP1-TKS (epPCR 1).** (**A**) Fluctuation assay results for ~90 colonies isolated from OR-Y488 transformed with either the epPCR 1 library (blue/orange) or the parent TP-DNAP1(I777K, L900S) (grey), then subject to URA3 selection. Note that per base mutation rate cannot be calculated by fluctuation analysis without copy number measurement; thus only per-cell mutation frequency is shown. Twelve library members (orange) were chosen for Sanger sequencing, using both mutation frequency and lineage (to minimize duplicate TP-DNAP1 variants) as criteria. Grey dotted line, mean mutation frequency for the three parent controls. (**B**) Mutation rate measurements for TP-DNAP1(I777K, L900S) and five unique TP-DNAP1 variants isolated from this directed evolution round. Following selection for URA3 and either fluctuation analysis (using trp5) or trp5 selection, individual clones were grown in -uracil growth medium for mutation accumulation on trp5. Two timepoints separated by 100 generations of growth were used as template for long-read high-throughput sequencing. Mutation rates are shown as a box plot and points for individual replicates where n > 1 biological replicate, or one line where n = 1. (**C**) Cumulative hamming distance distribution for all high throughput sequencing samples at the end of mutation accumulation.

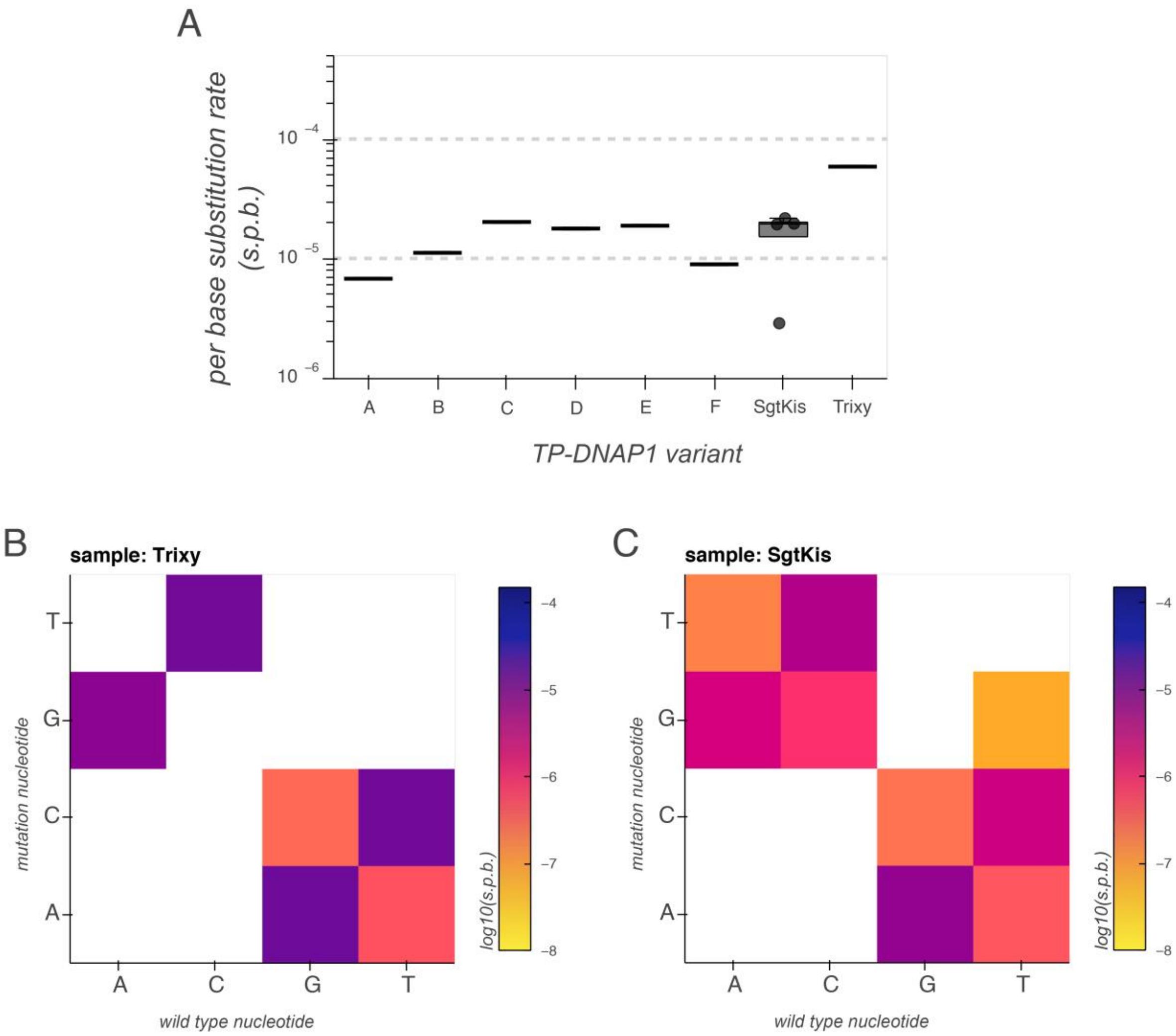


**Fig. S8. Identification of Trixy and SgtKis (epPCR 2).** (**A**) Mutation rate measurements for all substitutions from HTS dataset 3 by mutation accumulation. Mutation rates are shown as a box plot and points for individual replicates where n > 1 biological replicate, or one line where n = 1. (**B**-**C**) Heatmap representation of individual mutation rates for Trixy (B) and SgtKis (C) as measured in HTS dataset 3.

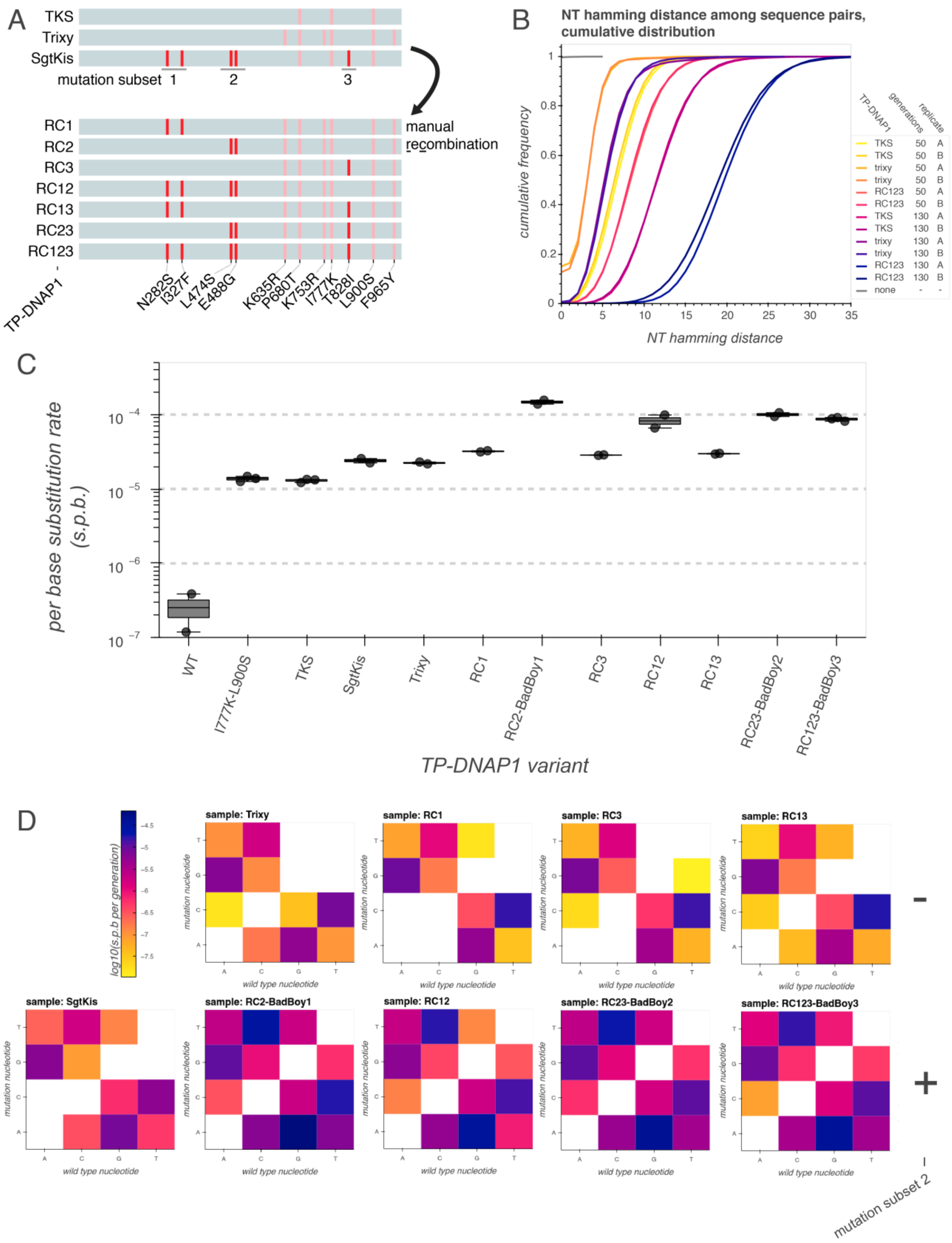


**Fig. S9. Mutation accumulation and HTS analysis of TP-DNAP1 variants from manual recombination.** (**A**) Illustration of manually constructed TP-DNAP1 variants derived from mutation subsets of SgtKis transplanted onto Trixy. (**B-D**) HTS analysis from an 80 generation mutation accumulation on a ~1 kb region of p1, highlighting the cumulative pairwise hamming distance for a subset of polymerase variants tested (B), the overall mutation rate measurements for all polymerases tested (C), and the effect of mutation subset 2 on the mutation spectrum by log transformed heatmap of individual mutation rates (D). Mutation rates in (B) are plotted as points for rates measured for each replicate and box plots of summary statistics for all biological replicates (n ≥ 2).

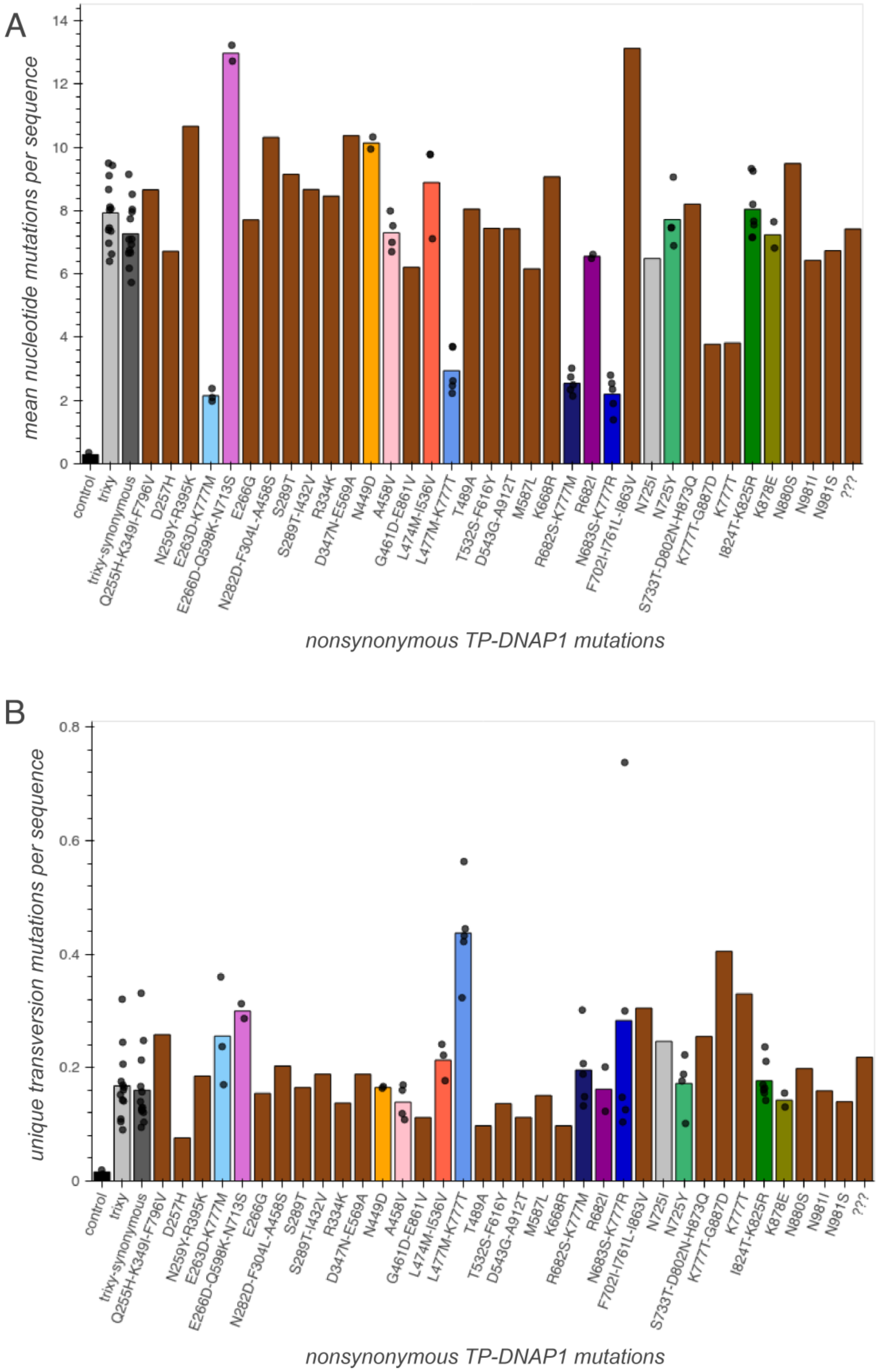
A
mean nucleotide mutations per sequence
nonsynonymous TP-DNAP1 mutations
B
unique transversion mutations per sequence
nonsynonymous TP-DNAP1 mutations
control
trixy
trixy-synonymous
Q255H-K349I-F796V
D257H
N259Y-R395K
E263D-K777M
E266D-Q598K-N713S
E266G
N282D-F304L-A458S
S289T
S289T-I432V
R334K
D347N-E569A
N449D
A458V
G461D-E861V
L474M-I536V
L477M-K777T
T489A
T532S-F616Y
D543G-A912T
M587L
K668R
R682S-K777M
R682I
N683S-K777R
F702I-I761L-I863V
N725I
N725Y
S733T-D802N-H873Q
K777T-G887D
K777T
I824T-K825R
K878E
N880S
N981I
N981S
???

**Fig. S10. Single timepoint HTS analysis of epPCR 3.** (**A-B**) An error prone PCR library (TP-DNAP1-Trixy template) was subject to either simultaneous or sequential selection for ura3* and trp5* reversion and a ~2kb region of p1 was PCR amplified and subject to HTS. Analysis of mean nucleotide (NT) mutations per sequence (A) and unique transversions per sequence (total unique transversion mutations / number of sequences analyzed) (B) was used to nominate mutations for the following round of directed evolution. Nonsynonymous amino acid substitutions in addition to those found in trixy are listed. Grey, no nonsynonymous TP-DNAP1 mutations in addition to tTP-DNAP1-Trixy mutations identified. Colors, multiple isolates contained the same polymerase. Brown, isolate contained a unique mutation combination. Blues, contains a mutation to residue 777.

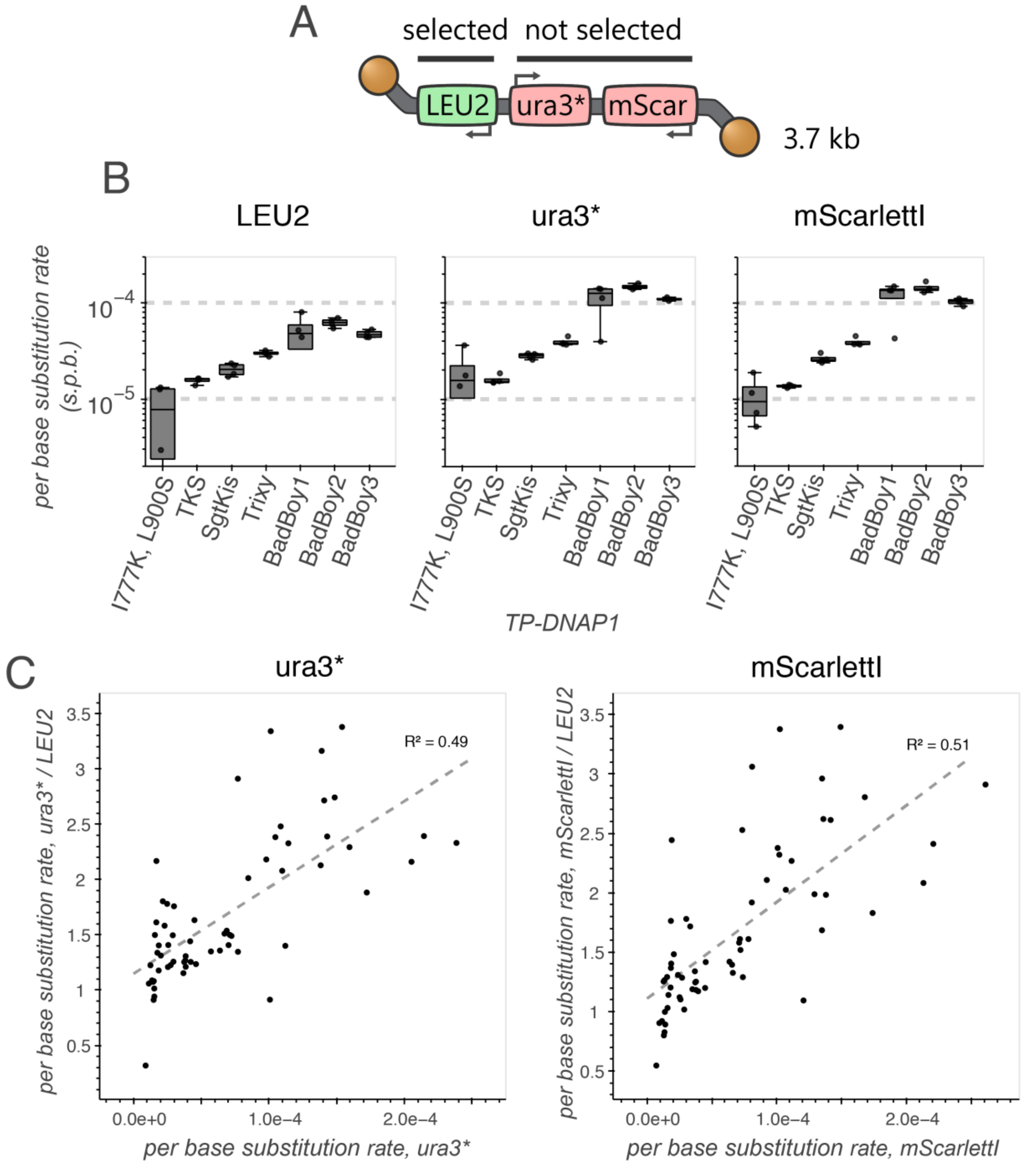


**Fig. S11. The effect of selection on mutation rate.** (**A**) Plasmid map for the p1 subjected to mutation accumulation in HTS dataset 6. Cells were passaged in +uracil/-leucine synthetic complete media resulting in functional selection for LEU2, but not ura3. (**B-C**) Comparison of mutation rates (HTS dataset 6) for regions of similar length (~1 kb), either under selection (LEU2) or not under selection (ura3*, mScarlettI) showing substitution rates for all substitution types as box plots and points (n=4 biological replicates) for a subset of TP-DNAP1 variants (B) or as a scatter plot (C) showing the relationship between mutation rate and fold change in mutation rate without (ura3* or mScarlettI) vs. with (LEU2) selection (one point per each n=1 individual polymerase variant replicate for all 17 TP-DNAP1 variants assayed).

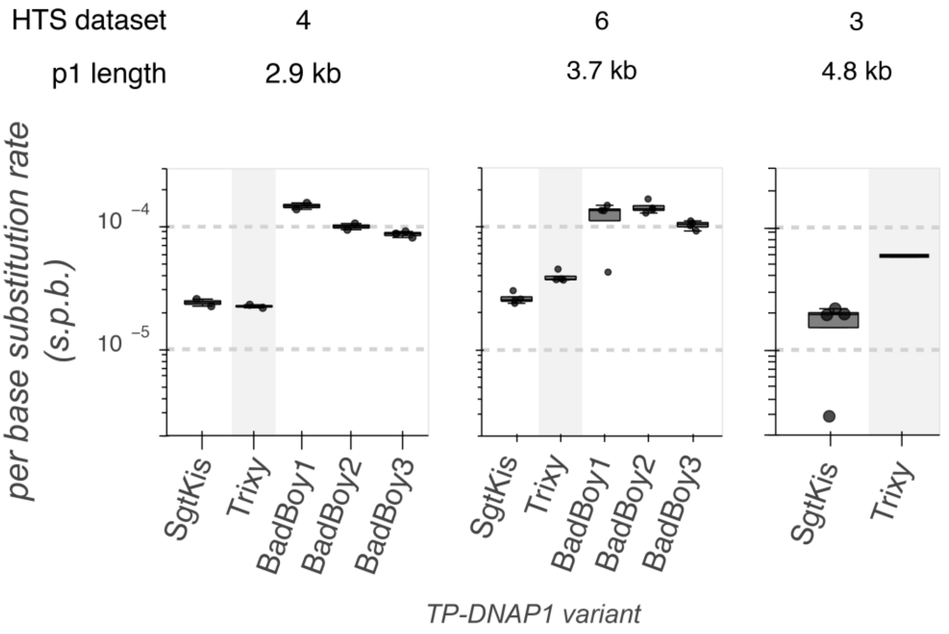


**Fig. S12. Relationship between p1 length and mutation rate.** Mutation rate measurements for a subset of TP-DNAP1 variants and the length of recombinant p1 used to generate the indicated mutation accumulation dataset are shown. TP-DNAP1-Trixy, whose mutation rates show the most obvious relationship with p1 length, is highlighted in grey. Data are identical to those found in Figs. S9, 1C, and S8 (left to right).

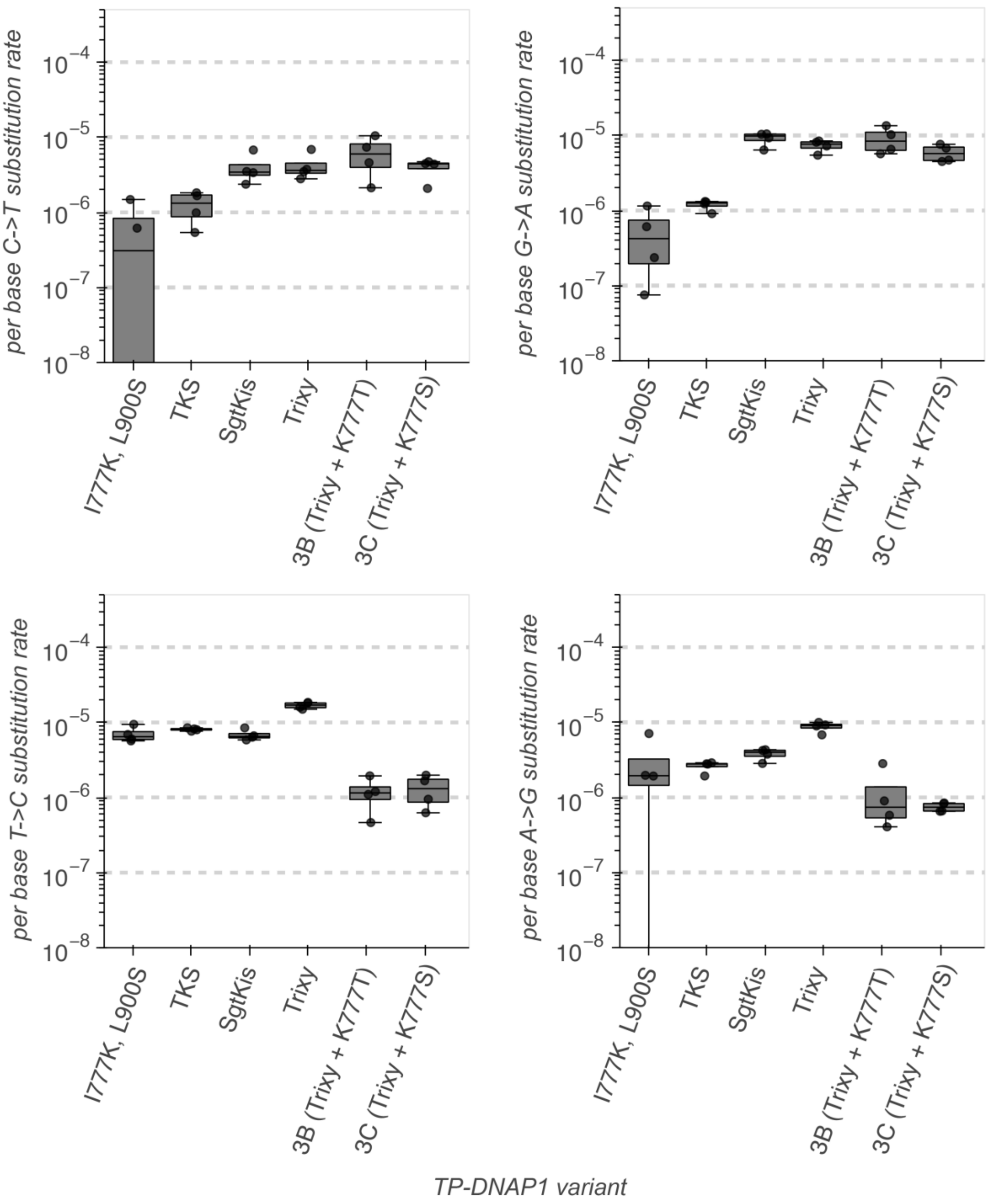


**Fig. S13. The effect of mutations to residue 777 on transition mutation rates.** Mutation rate measurements from HTS dataset 6 for all TP-DNAP1 variants assayed for all four transition mutations, highlighting variants 3B, 3C, which differ from Trixy only by a single mutation (K777T/K777S, respectively) and BB-3B and BB-3C, which differ from BadBoy3 by only a single mutation (K777T/K777S, respectively).

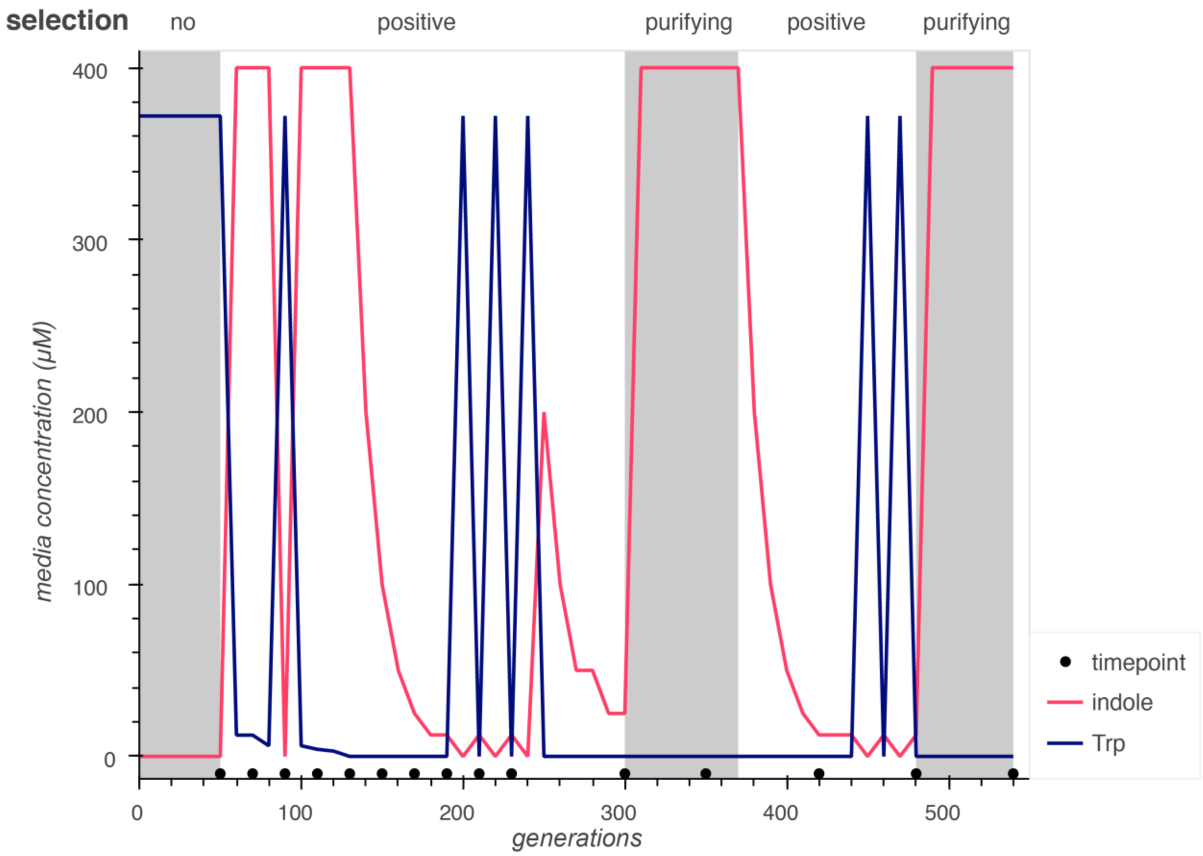


**Fig. S14. Selection condition schedule for TrpB evolution.** All 96 cultures in the experiment were passaged in synthetic complete medium with the indicated concentrations of indole and Trp, and DNA was harvested and sequenced at the indicated timepoints. Intermittent passages into Trp-supplemented media during positive selection phases were employed to increase the rate of diversification.

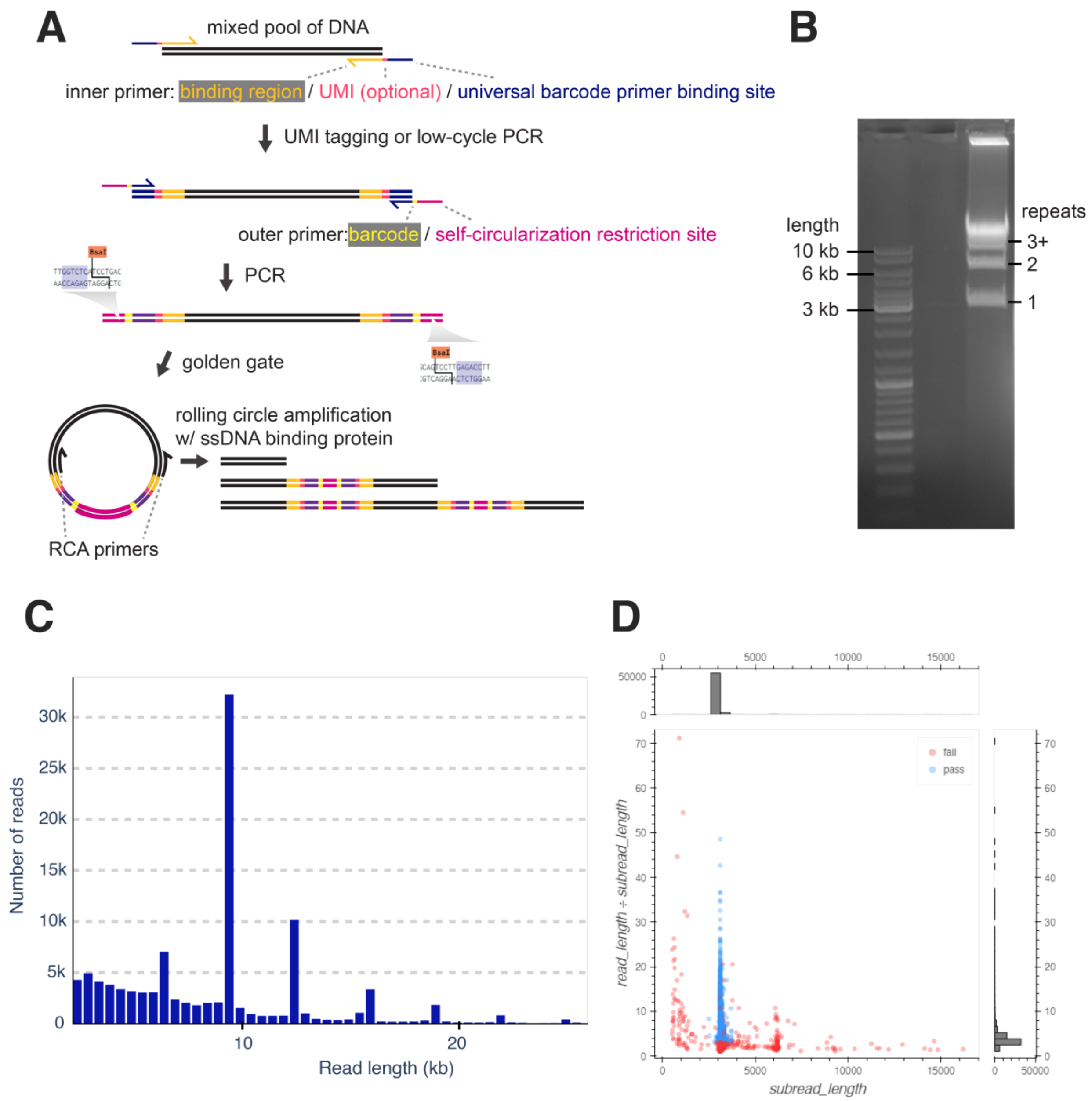


**Figure S15. A modified rolling circle amplification (RCA) method for high accuracy long read nanopore sequencing.** (**A**) Illustration of the steps involved in the RCA method. Note that RCA primers must bind internally to the inner primers to prevent amplification of primer dimers. (**B**-**D**) An example RCA and nanopore sequencing results using a Flongle flow cell for an amplicon of 3 kb in length, showing gel electrophoresis of the RCA product (B), the distribution of read lengths (C), and the distribution of the number of repeats (estimated as the ratio of read length to subread length) vs. subread length (D). Number of repeats and subread length were used for quality filtering prior to downstream analysis, the results of which are shown in color in D. Note that the bands corresponding to 3+ repeats were gel extracted for sequencing, resulting in the minimal amplicons with 2 or fewer repeats observed in C.

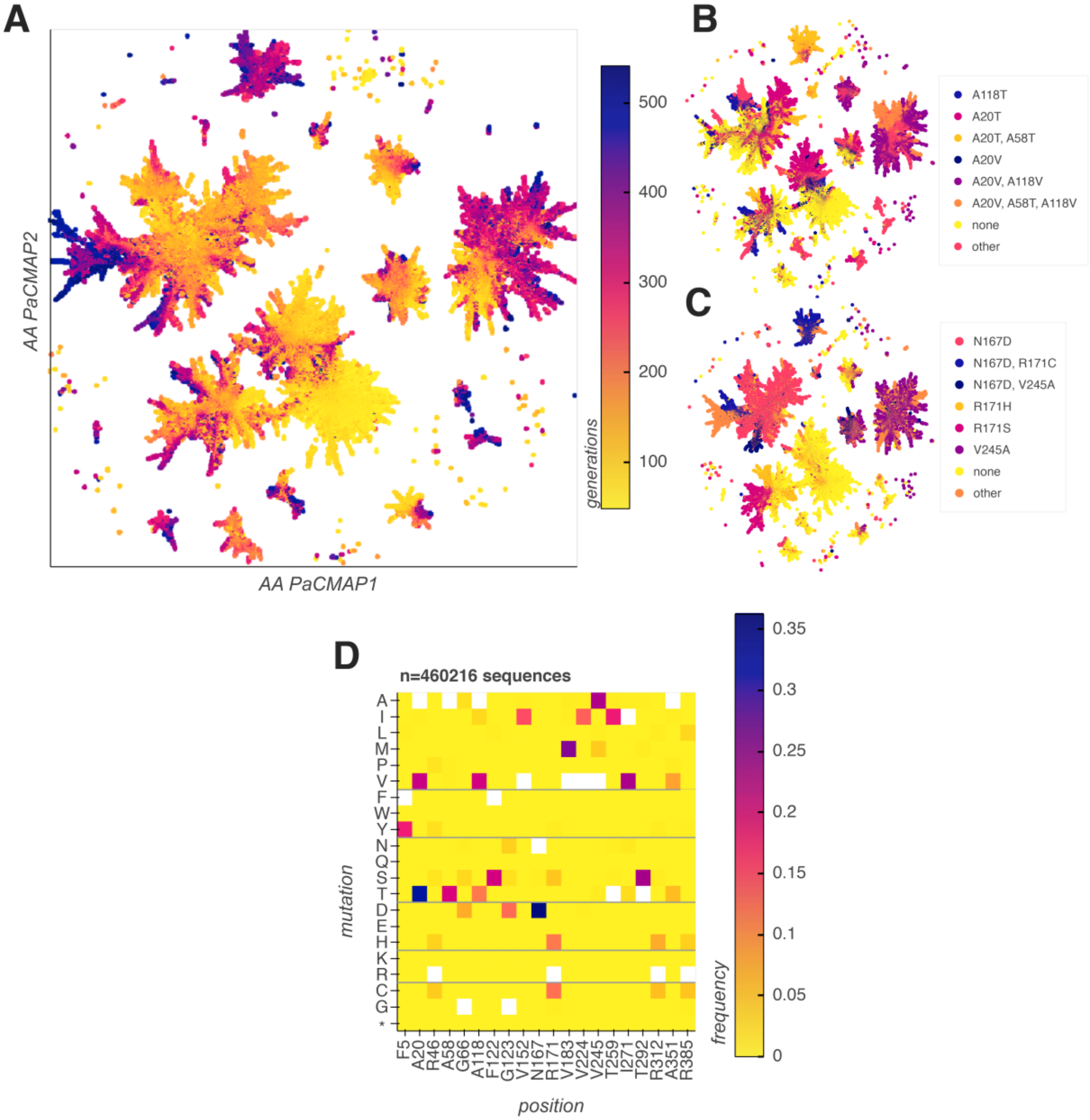


**Figure S16. Diversity of evolved TrpB variants.** (**A**) 2-dimensional representation for all unique genotypes identified using PaCMAP for dimensionality reduction, with the timepoint from which each genotype was identified represented as color. (**B** and **C**) 2D representations as in E, with color used to indicate the most frequently observed combinations of mutations among the six most frequently mutated positions in the final timepoint. (**D**) Heatmap of all mutations to the 20 most commonly mutated positions throughout the entire experiment.

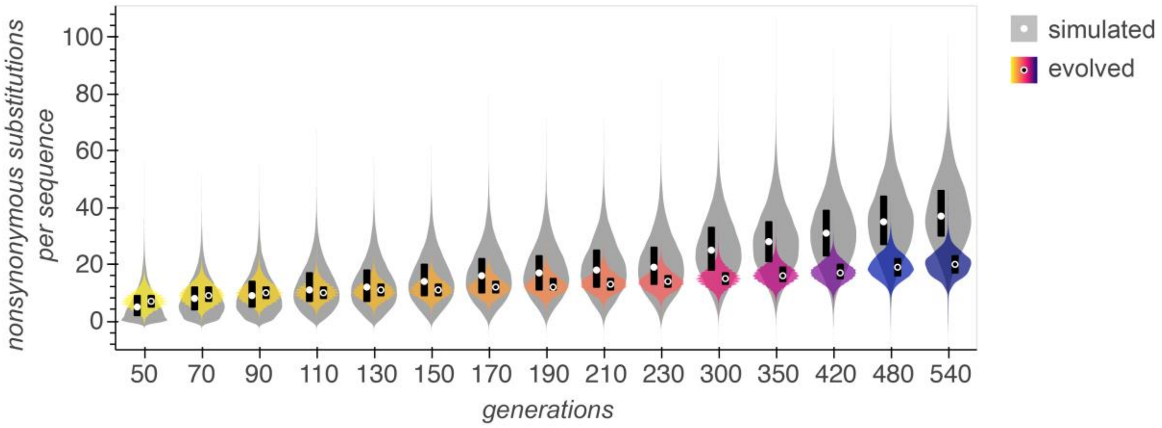


**Figure S17. Nonsynonymous mutation distributions for simulated and evolved sequences.** Violin plots of nonsynonymous mutations per sequence in each timepoint of TrpB evolution (colors) compared to that of sequences from a simulated dataset with an equivalent number of synonymous mutations generated at random using mutation rates and preferences of BadBoy2 (grey). Points and black bars denote the means and interquartile range for all sequences within each timepoint.

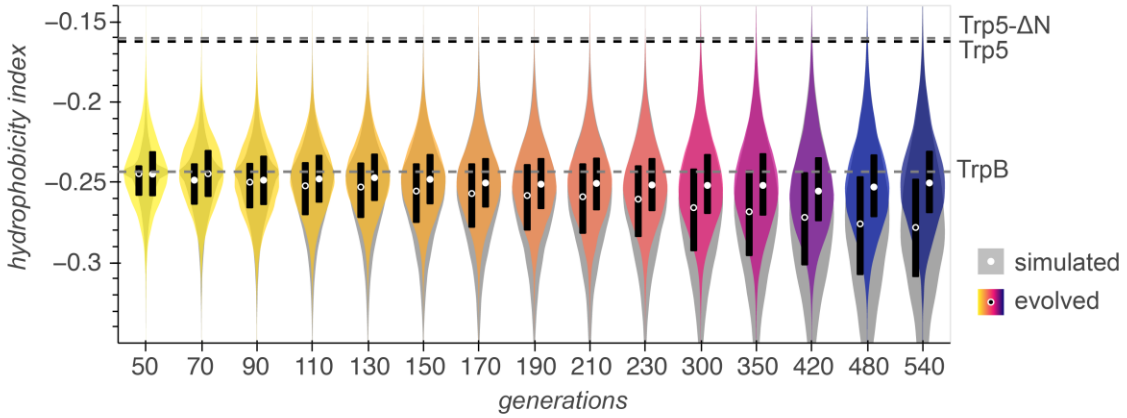


**Figure S18. Effect of long-term mutagenesis on hydrophobicity.** Violin plots of Kyte-Doolittle hydrophobicity index in each timepoint of TrpB evolution for both evolved (colors) and simulated sequences (grey). Points and black bars denote the means and interquartile range for all sequences within each timepoint. Hydrophobicity indices of the wild type TrpB, the TrpA-TrpB holoenzyme ortholog from *Saccharomyces cerevisiae* (Trp5) and an N-terminally-truncated Trp5 homologous to TrpB (Trp5-ΔN) are shown for comparison.

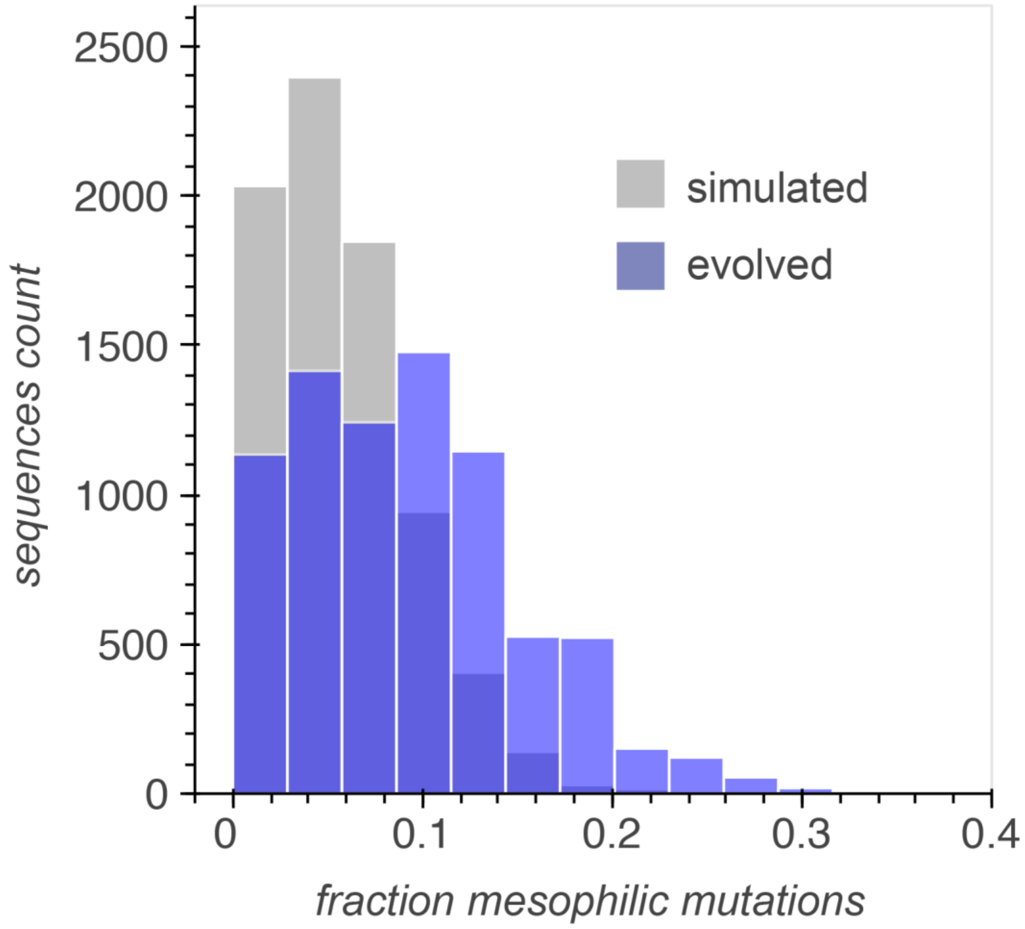


**Figure S19. Distribution of mesophile mutation fraction for both evolved and simulated TrpB sequences.** Mesophile mutation fraction was calculated for each sequence as the number of mesophile mutations divided by the total mutations. A set of 17 amino acid replacements (*e.g.* Proline to Serine) identified by Haney *et al.*[14] were designated as mesophile mutations (see Methods). Only sequences from the final timepoint (540 generations) were included.

### References


1. Rix, G. *et al.* Scalable continuous evolution for the generation of diverse enzyme variants encompassing promiscuous activities. *Nat. Commun.* **11**, 1–11 (2020).

2. Alani, E., Cao, L. & Kleckner, N. A method for gene disruption that allows repeated use of URA3 selection in the construction of multiply disrupted yeast strains. *Genetics* **116**, 541–545 (1987).

3. Ryan, O. W. & Cate, J. H. D. *Multiplex engineering of industrial yeast genomes using CRISPRm*. *Methods in Enzymology* vol. 546 (Elsevier Inc., 2014).

4. Zurek, P. J., Knyphausen, P., Neufeld, K., Pushpanath, A. & Hollfelder, F. UMI-linked consensus sequencing enables phylogenetic analysis of directed evolution. *Nat. Commun.* **11**, 1–10 (2020).

5. Volden, R. *et al.* Improving nanopore read accuracy with the R2C2 method enables the sequencing of highly multiplexed full-length single-cell cDNA. *Proc. Natl. Acad. Sci. U. S. A.* **115**, 9726–9731 (2018).

6. Oliynyk, R. T. & Church, G. M. Efficient modification and preparation of circular DNA for expression in cell culture. *Commun. Biol.* **5**, 1–10 (2022).

7. Zhang, Y. & Tanner, N. A. Isothermal Amplification of Long, Discrete DNA Fragments Facilitated by Single-Stranded Binding Protein. *Sci. Rep.* **7**, 1–9 (2017).

8. Ravikumar, A., Arzumanyan, G. A., Obadi, M. K. A., Javanpour, A. A. & Liu, C. C. Scalable, Continuous Evolution of Genes at Mutation Rates above Genomic Error Thresholds. *Cell.* **175**, 1946-1957.e13 (2018).

9. Rubin, A. F. *et al.* A statistical framework for analyzing deep mutational scanning data. *Genome Biol.* **18**, 1–15 (2017).

10. Notin, P. *et al.* TranceptEVE: Combining Family-specific and Family-agnostic Models of Protein Sequences for Improved Fitness Prediction. *bioRxiv* 2022.12.07.519495 (2022).

11. Johnson, L. S., Eddy, S. R. & Portugaly, E. Johnson, L.S.; Eddy, S.R.; Portugaly, E. Hidden Markov model speed heuristic and iterative HMM search procedure. BMC Bioinform. 2010, 11, 431. *BMC Bioinformatics* **11**, 431 (2010).

12. Hopf, T. A. *et al.* Mutation effects predicted from sequence co-variation. *Nat. Biotechnol.* **35**, 128–135 (2017).

13. Cock, P. J. A. *et al.* Biopython: Freely available Python tools for computational molecular biology and bioinformatics. *Bioinformatics* **25**, 1422–1423 (2009).

14. Haney, P. J. *et al.* Thermal adaptation analyzed by comparison of protein sequences from mesophilic and extremely thermophilic Methanococcus species. *Proc. Natl. Acad. Sci. U. S. A.* **96**, 3578–3583 (1999).

15. Lee, M. E., DeLoache, W. C., Cervantes, B. & Dueber, J. E. A Highly Characterized Yeast Toolkit for Modular, Multipart Assembly. *ACS Synth. Biol.* **4**, 975–986 (2015).
